# Colonization at birth with human CST IV cervicovaginal microbiota alters development and increases neonatal mortality in mice

**DOI:** 10.1101/2021.02.01.429213

**Authors:** Eldin Jašarević, Elizabeth M. Hill, Patrick J. Kane, Lindsay Rutt, Trevonn Gyles, Lillian Folts, Kylie D. Rock, Christopher D. Howard, Kathleen E. Morrison, Jacques Ravel, Tracy L. Bale

## Abstract

Profound racial health disparities contribute to maternal-infant morbidity and mortality. An emergent risk factor is the maternal microbiota, whereby compositional alterations impact maternal health and offspring development during pregnancy and beyond. The presence of a nonoptimal CST IV cervicovaginal microbiota, more common in Black and Hispanic women, is associated with increased risk of preterm birth and adverse birth outcomes. Through examination of the biological mechanisms by which vertical transmission of microbiota from mother to offspring influences postnatal development, we found that exposing cesarean delivered mice with CST IV cervicovaginal microbiota from pregnant women produced lasting effects on offspring metabolic, immune, and neural outcomes. We then examined how compounding effects of a typical high-risk, proinflammatory *in utero* environment, characterized by a maternal obesogenic state and the presence of *G. vaginalis*, would affect the offspring response to CST IV microbial gut colonization. The resultant developmental immaturity, coupled with an exaggerated immune response induced by exposure to risk-associated maternal microbiota, resulted in a profound increase in neonatal mortality, supporting the critical importance of elucidating the multifactorial biological mechanisms involved in high-risk pregnancies.

**Highlights:**

- Gut colonization by nonoptimal human cervicovaginal microbiota had sex-specific effects on postnatal development.
- A high-risk *in utero* environment increased offspring mortality risk.
- Heightened innate immune response associated with mortality risk.
- Developmental immaturity, premature birth, and exposure to CST IV contribute to increased offspring mortality risk.

## Introduction

Profound racial and ethnic disparities exist in maternal-child health outcomes (Bryant et al., 2010; Hauck et al., 2011; Hebert et al., 2008; Howell, 2018; Lu and Halfon, 2003; MacDorman and Mathews, 2011; Mehra et al., 2019; Orr et al., 2012; Osypuk and Acevedo-Garcia, 2008). Black women in the United States are four times more likely to experience preterm birth, fetal growth restriction, and maternal and infant morbidity and mortality than white women (Balchin and Steer, 2007; Chinn et al., 2020; Tangel et al., 2019; Tucker et al., 2007; Wollmann, 1998). These nationwide trends likely underestimate the widening gap in racial disparities in maternal-child health outcomes that has been reported at the state, county, and city-level (March of Dimes Premature Birth Report Card; (Frey and Klebanoff, 2016; Gyamfi-Bannerman and Ananth, 2014; Tangel et al., 2019). For instance, the preterm birth rate among Black women in Baltimore City is 44% higher than the rate among all other women (i.e., a 13% preterm birth rate among Black women compared with 8.8% among white women in Baltimore City; National Center for Health Statistics). Collectively, these patterns highlight a critical need for elucidation into the biological mechanisms underlying these increased risks.

Among these well-documented social and environmental determinants of racial health disparities, the cervicovaginal microbiota has emerged as an important biological contributor to maternal and offspring health outcomes (Aagaard et al., 2012; Brown et al., 2019; Elovitz et al., 2019; Fettweis et al., 2019; Florova et al., 2021; Gajer et al., 2012; Lorch and Enlow, 2016; Owens and Fett, 2019; Ravel et al., 2011; Romero et al., 2014; Serrano et al., 2019; Williams and Sternthal, 2010). In contrast to the gut microbiota, an optimal cervicovaginal microbiome consists of minimally diverse microbial communities that recent cultivation-independent approaches have classified into five primary community state types (CST) (France et al., 2020; Ravel et al., 2011). Generally, CST I, II, III and V are dominated by the presence of *Lactobacillus* species while the nonoptimal cervicovaginal microbiome, CST IV, is defined by an absence of *Lactobacillus* and the presence of a wide array of anaerobic bacteria, including *Gardnerella vaginalis* and *Atopobium vaginae* (Fredricks et al., 2005; Gajer et al., 2012; Greenbaum et al., 2019; redondo-Lopez et al., 1990; Ravel et al., 2011; Romero et al., 2014). During pregnancy, Black women are more likely to harbor CST IV microbiota that have been associated with increased incidence of obstetric complications, inflammation in the intrauterine environment, spontaneous preterm birth, and neonatal morbidity (Brown et al., 2018, 2018; Callahan et al., 2017; DiGiulio et al., 2015; Elovitz et al., 2019; Fettweis et al., 2019; Florova et al., 2021; Gerson et al., 2020a; Hočevar et al., 2019; Ravel et al., 2011; Romero et al., 2014; Tabatabaei et al., 2019, 2019; Zhou et al., 2007). The risk associated with CST IV may be further amplified by a variety of environmental factors that promote chronic and systemic inflammation, including maternal stress and trauma, infection, and consumption of a high-fat, low-fiber diet (Brookheart et al., 2019; Danese and Lewis, 2017; Gerson et al., 2020b; Shivakoti et al., 2020).

In addition to the importance of the cervicovaginal microbiome with respect to maternal health, the cervicovaginal microbiota is an important microbial reservoir to which the newborn is exposed during a vaginal birth (Bokulich et al., 2016; Dominguez-Bello et al., 2010, 2016). Recent strain-resolved analyses have shown that maternal cervicovaginal bacteria, such as *L. crispatus*, *G. vaginalis,* and *A. vaginae,* are among the earliest colonizers to persist within the infant intestinal tract during the first few days after birth (Asnicar et al., 2017; Ferretti et al., 2018; Gabriel et al., 2018; Yassour et al., 2018). Preclinical animal models have shown that pioneer communities, such as those from the maternal cervicovaginal and gut microbiota, colonize the mucosal surfaces of the newborn and initiate processes that include immune system maturation, nutritional provisioning for the developing brain, and other adaptations to the postnatal environment (Arrieta et al., 2015; Fulde et al., 2018; Gensollen et al., 2016; Jašarević and Bale, 2019; McDonald and McCoy, 2019; Renz et al., 2018; Walker, 2017). Indeed, these animal studies have shown that disruption in this intestinal colonization process promotes lasting changes to the offspring’s immune system, hypothalamic-pituitary-adrenal stress axis, as well as increased susceptibility to allergy, obesity, and diabetes across the lifespan (Al Nabhani et al., 2019a, 2019b; Arrieta et al., 2015; Jašarević et al., 2015, 2017, 2018; Morais et al., 2020; Sudo et al., 2004). While such studies support the notion that vertical transmission of maternal microbiota impacts offspring development in a composition-dependent manner, the specific health outcomes following colonization by a nonoptimal CST IV cervicovaginal microbiome, especially in the context of a high-risk pregnancy, have not been examined.

To begin addressing these complex interactions, we first established a protocol that simulates unique features of early life colonization in humans by oral gavaging cesarean delivered mice with distinct human maternal microbiota, comparing transplants from optimal, *Lactobacillus crispatus* dominant, CST I microbiota with nonoptimal cervicovaginal microbiome (CST IV). We assessed the impact of these communities on postnatal gut and immune development, growth and metabolism, circulating immunity, and neural circuits involved in the control of feeding and metabolism in these mice. Moreover, to then model a high-risk *in utero* environment, we added the compounding effects of maternal obesity, diabetes, and a presence of *G. vaginalis* that is common to CST IV women, to ascertain the additive impact on the offspring response to CST IV microbial colonization at birth.

## Results

### Establishment of a mouse model to determine the functional attributes of the human maternal microbiota on offspring outcomes

We first sought to establish a preclinical mouse model that may capture the complex metabolic, immune and neurodevelopmental consequences of early-life colonization by defined communities of the human maternal microbiota. Given the recent public and scientific interest in reconstituting the early-life microbiome of cesarean delivered infants through means of transplanting these newborns with maternal fecal or cervicovaginal microbiota, preclinical models that may provide mechanistic insight into the metabolic and immunological consequences of these communities is paramount (Dominguez-Bello et al., 2016; Korpela et al., 2020; Limaye and Ratner, 2020; Mortensen et al., 2021; Mueller et al., 2020). To determine whether transplantation of human cervicovaginal microbiota to newborns exert lasting effects, we used our C-section and oral gavage method to manipulate the microbial communities that colonize the newborn intestinal tract of conventionally-reared mice and assessed lasting outcomes (Jašarević et al., 2018). Late-pregnancy samples (gestational week 36 - 39) were selected from a large prospectively-collected cohort of pregnant women in order to capture the microbial communities with the greatest likelihood that may be transferred to newborns during birth. While there are numerous microbial communities transferred from mother to newborn, we limited our experiments to CST I and CST IV microbiota samples as members of these microbial community have been detected in the stool of infants (Ferretti et al., 2018). Additionally, CST IV and CST I microbiota exhibit divergent functional, metabolic, and immunological effects on the maternal cervicovaginal space (Smith and Ravel, 2017), suggesting that vertical transmission of these microbiota may influence offspring development in a CST-dependent manner.

To examine this hypothesis, on embryonic day (E) 18.5 C57BL/6J mouse pups were delivered by cesarean and colonized with human cervicovaginal CST I or CST IV microbiota by oral gavage (Figure 1A). To control for effects of inoculation, we included a control group in which C-section delivered pups were inoculated with the Amies transport medium, in which the human cervicovaginal swab samples were stored. Age-matched vaginally delivered (VD) offspring were also included to assess the lasting effects of mode of delivery, early birth (i.e., the C-section offspring being born 0.5 days early), and to account for effects of being colonized by human- and mouse-specific microbiota. Further, we controlled for confounding effects of the postnatal environment and housing conditions by cross-fostering both sexes and all treatment groups to surrogate 129/SvJ foster dams. The 129/SvJ strain was chosen due to the well-documented occurrence of infanticide and rejection of non-genetically related offspring in the C57BL/6J strain, but not 129/SvJ (Crawley et al., 1997). This modular approach was able to delineate the unique contributions of offspring genetics, maternal genetics, human cervicovaginal microbiota, and the postnatal environment, as C-section pups inoculated with CST I and CST IV share the same strain background and postnatal environment but are colonized with human cervicovaginal microbiota that differ in composition and expressed functions.

**Figure 1.**
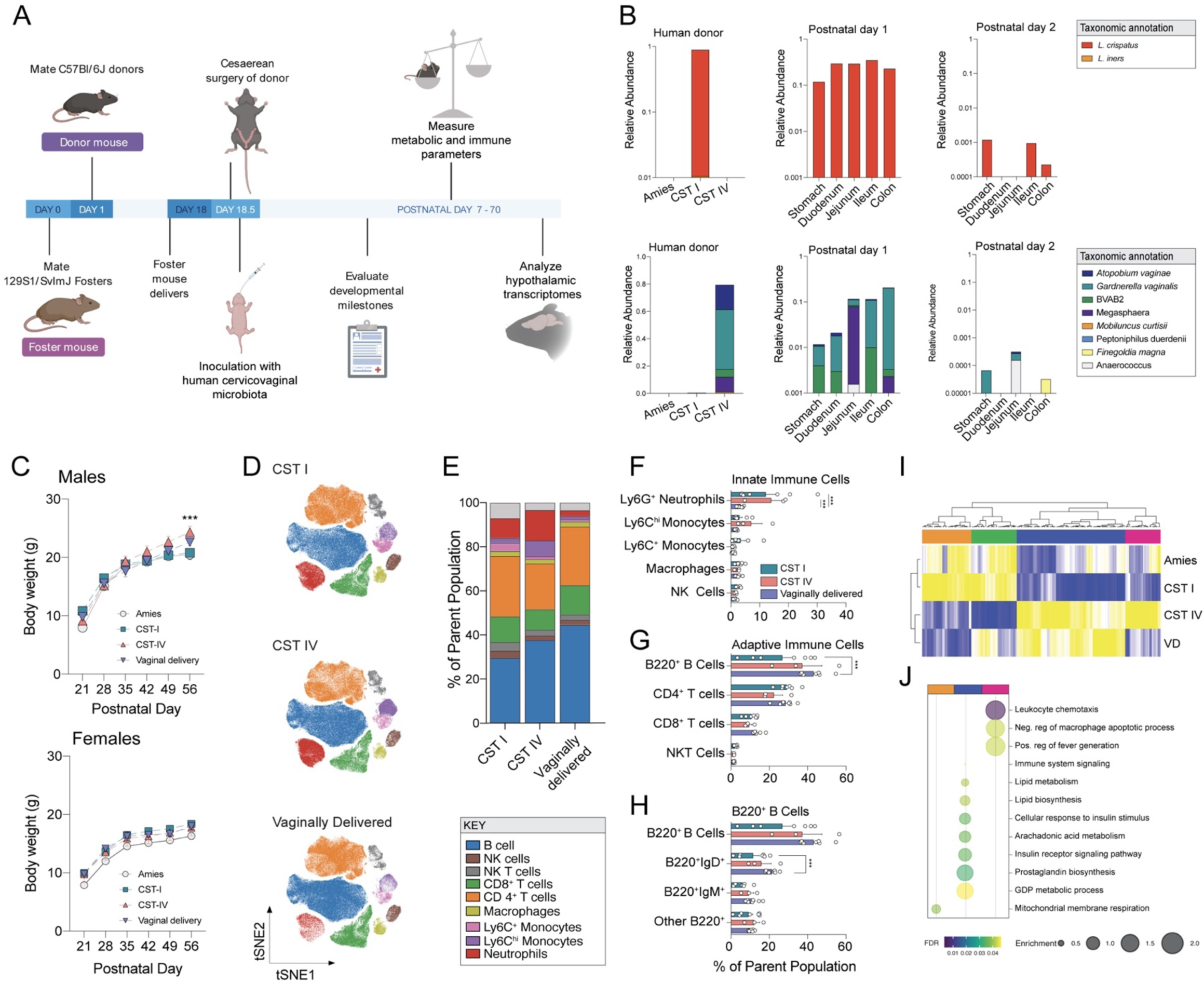
Early life exposure to human cervicovaginal microbiota results in lasting effects on growth, circulating immunity, and hypothalamic gene expression patterns. **(A)** Schematic of the experimental timeline and protocol used to establish a mouse model to investigate the impact of exposure to maternal human cervicovaginal microbiota at birth on lasting offspring outcomes. See *Methods* for details. **(B)** 16S rRNA-based validation of presence of human vaginal microbiota across intestinal segments of C-section neonate mice inoculated with *L. crispatus-*dominated microbiota (CST I) or lactobacilli-deficient, nonoptimal microbiota (CST IV) collected from late-gestation women. *Upper left panel*, mean relative abundance of *L. crispatus* showing no detectable amplification of *L. crispatus* in the Amies transport medium and CST IV samples. *Upper middle and right panel*, mean relative abundance of *L. crispatus* in CST I inoculated offspring at postnatal day 1 and 2. No amplification of *L. crispatus* was detected in Amies and CST IV inoculated pups at either timepoint. *Lower left panel*, mean relative abundance from CST IV associated taxa showing no detectable amplification in the Amies transport medium and CST I samples. *Upper middle and right panel*, mean relative abundance of CST IV inoculated offspring at postnatal day 1 and 2. No amplification of CST IV was detected in Amies and CST I inoculated pups at either timepoint. N = 7 – 20 per group and intestinal segment. **(C)** Sex-specific effects of CST I and CST IV inoculation on body weight across development. *Top panel*, male body weight was significantly changed over time (two-way ANOVA, main effect of time F_5, 170_ = 689.4, *P* < 0.0001) and the interaction between time and treatment (two-way ANOVA, time*treatment interaction, F_15, 170_ = 8.544, *P* < 0.0001). Tukey’s post hoc analysis revealed that CST IV males weighed more than Amies and CST I males at P56 (Tukey’s, *P* = 0.0009 and 0.0006, respectively). No differences in body weight in CST I males compared with Amies or vaginally delivered males at P56 (Tukey’s, *P* = 0.990 and *P* = 0.261). *Bottom panel,* female body weight was significantly changed over time (two-way ANOVA, main effect of time F_5, 130_ = 306.819, *P* < 0.0001). No effect of treatment or their interaction was observed. N = 7 – 12 animals per sex, treatment, and timepoint. Data represented as mean ± SEM. Data is representative of two independent experiments. *** *P <* 0.001. **(D-H)** High-dimensional, single-cell mapping reveals lasting effects of CST I and CST IV inoculation on the circulating immune compartment in adult males. **(D)** *t-*Distributed stochastic neighbor embedding (t-SNE) visualization demonstrating CyTOF phenotyping of CD45+ immune cells in whole blood of CST I, CST IV, and vaginally delivered postnatal day 56 adult males (equal sampling across treatment groups, total 390,000 events). **(E)** Average frequencies of major leukocytes within whole blood in CST I, CST IV, and VD adult males. **(F)** Frequencies of circulating innate immune cells showing increased frequency of neutrophils in CST I and CST IV males compared with VD males at P56 (two-way ANOVA, treatment*immune cell interaction, F_8, 56_ = 2.870, P = 0.0095; Tukey’s post-hoc CST I vs VD *P* = 0.0001; Tukey’s post-hoc CST IV vs VD *P* = 0.0003). N = 3 - 7 males per treatment. Data represented as mean ± SEM with individual data points overlaid. *** *P <* 0.001 **(G)** Frequencies of circulating adaptive immune cells showing decreased frequency of B220+ B cells in CST I males compared with VD males at P56 (two-way ANOVA, treatment*immune cell interaction, F_6, 56_ = 2.492, *P* = 0.0330; Tukey’s post-hoc CST I vs. VD *P* = 0.0003). N = 3 −7 males per treatment. Data represented as mean ± SEM with individual data points overlaid. *** *P <* 0.001 **(H)** Frequencies of circulating B220^+^ B cells showing decreased frequency of B220^+^IgD^+^ B cells in CST I males compared with VD males (two-way ANOVA, treatment*immune cell interaction, F_4, 28_ = 5.471, P = 0.0022; Tukey’s post-hoc CST I vs. VD *P* = 0.0017). *** *P <* 0.001 **(I)** Heatmap depicting mean expression of genes in the paraventricular nucleus of the hypothalamus in P56 males. Paralleling body weight differences between CST I and CST IV males at P56, RNAseq analysis shows differences in PVN gene expression patterns between CST I and CST IV males (linear fit model, FDR < 0.1, log(FC) = 1.5). Unbiased hierarchical clustering showing similarity in transcriptional patterns between CST IV and VD males, and CST I and Amies males. *Z*-scores plotted across individuals for each gene. N = 4 – 6 males per treatment. **(J)** Cluster-based functional enrichment analysis of differentially expressed genes in the paraventricular nucleus of the hypothalamus in P56 males. Significant enrichment of pathways involved in metabolism and immunity in CST IV compared with CST I males, showing unique pathways in the PVN that are associated with body weight changes in CST IV males (FDR < 0.05). Bubble plot size denotes enrichment. N = 4 – 6 males per treatment.

Following these guidelines, we next determined whether offspring gut microbiota composition is influenced by inoculation of human cervicovaginal microbiota using 16S ribosomal RNA (rRNA) gene sequence profiling to characterize and compare the composition of the gut microbiota from C-section pups inoculated with Amies, CST I or CST IV and age-matched VD pups at postnatal day 1, corresponding to 24 hours post-inoculation (Figure 1B). Differential abundance analysis revealed that no taxa from CST I or CST IV were detected in Amies inoculated pups. Similarly, CST I inoculated pups showed no taxa associated with CST IV microbiota, and CST IV inoculated pups showed no taxa associated with CST I microbiota. Using a complimentary qPCR method for analysis of total counts of the *rpoB* gene from *G. vaginalis* –a common member of CST IV– we confirmed significant increase in total *G. vaginalis* in CST IV compared with CST I, Amies, or VD pups (Supporting Figure 1). Consistent with previous reports showing transient colonization by cervicovaginal microbiota, we observed a reduction in the recovery of human cervicovaginal microbiota from the intestinal tract of C-section delivered and inoculated mice by postnatal day 2, corresponding to 48 hours post-inoculation (Bennet et al., 1992; Ferretti et al., 2018) (Figure 1B).

To identify microbial reservoirs that may colonize neonatal mice at postnatal day 2, 16S rRNA gene sequence profiling was used to characterize the microbiota from environmental and maternal origin, including corn cob bedding, nestlet material, maternal skin surrounding mammary glands, mouth, gut, and vagina (Supporting Figure 2A). Non-metric multidimensional scaling (NMDS) analysis of maternal and environmental samples revealed that environmental samples formed a unique cluster from maternal samples (Supporting Figure 2B). Comparison of taxa between maternal and environmental samples revealed site-specific enrichment of microbiota, including amplicon sequence variants that mapped to *Pasteurella pneumotropica* recovered from maternal skin and vaginal fluid samples, and *Actinobacillus muris* recovered from maternal skin surrounding the mammary gland (Supporting Figure 2C). Additional NMDS analysis of maternal, C-section offspring, and environmental samples revealed no overlap between environmental and C-section offspring, suggesting that the local environment is not a significant reservoir of microbiota colonizing C-section offspring at postnatal day 2 (Supporting Figure 2D). Conversely, samples from C-section offspring and maternal skin samples showed considerable overlap, and comparison of taxa across intestinal segments of Amies, CST I, and CST IV offspring revealed that *P. pneumotropica* is the most abundant taxa present in C-section offspring at postnatal day 2 (Supporting Figure 2E-H). Given the presence of *P. pneumotropica* and *A. muris* on maternal skin and confirmation of a milk dot in all offspring, this suggests that the maternal skin is an important reservoir for colonizing the C-section mouse pup intestine. To determine whether colonization by CST I and CST IV impacted offspring gut microbiota composition, we used linear discriminant analysis – effect size (LEfSe) to test for differential abundance across intestinal segments of CST I and CST IV pups at postnatal day 2. Differential abundance analysis revealed increased abundance of *Gammaproteobacteria* and *Escherichia-Shigella* in CST IV pups compared with CST I pups at postnatal day 2 (Supporting Figure 2I,J). No other taxa differed between CST I and CST IV pups. Taken together, our results demonstrating transient colonization of the neonatal mouse gut by human cervicovaginal microbiota are consistent with human studies assessing maternal-to-infant microbiota transmission from different maternal body sites (Ferretti et al., 2018).

**Figure 2.**
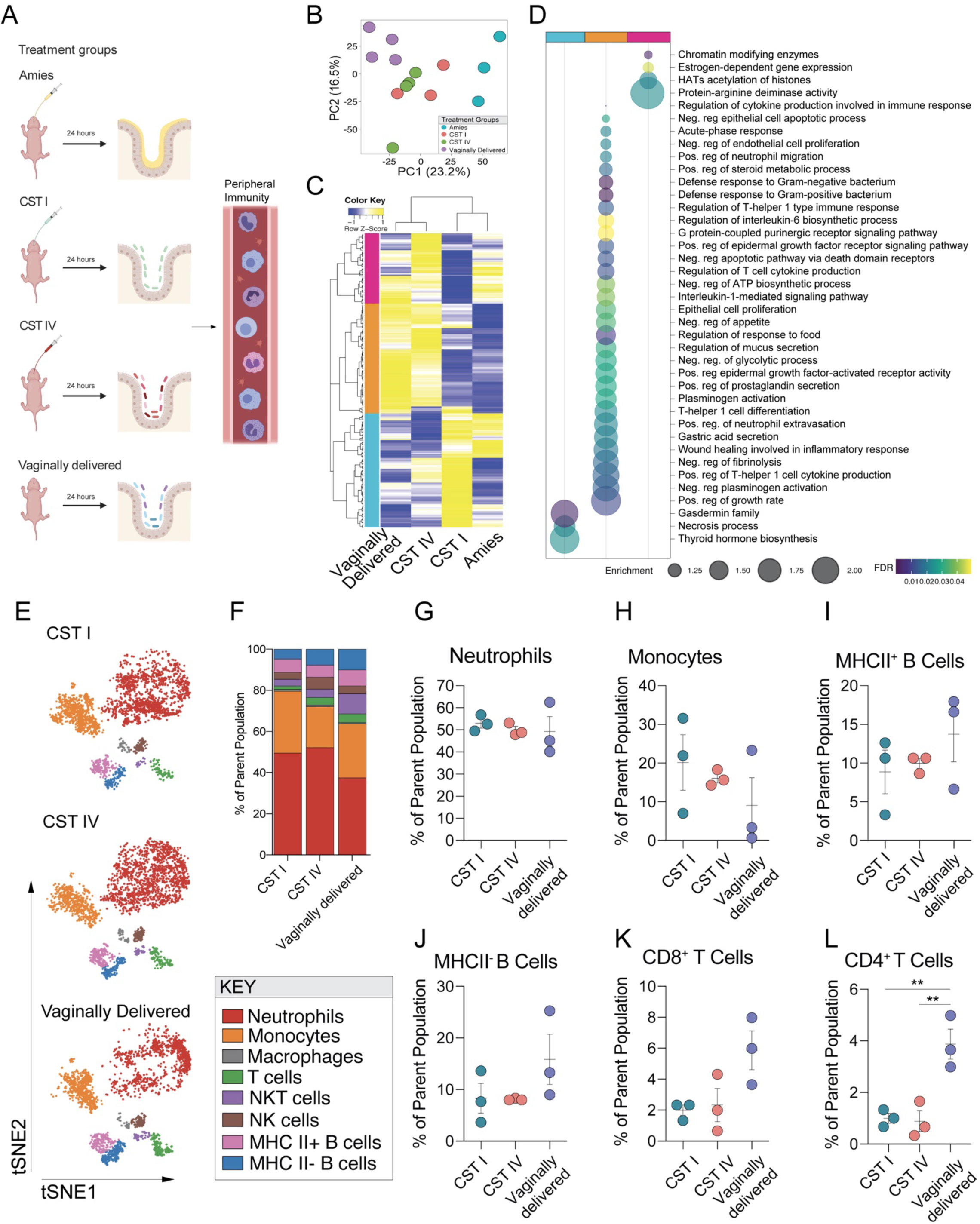
Exposure to human cervicovaginal microbiota effects transcriptional profiles of the neonatal ileum and contributes to the circulating immune compartment in a community-dependent manner. **(A)** Schematic of experimental outline to determine the influence of the human microbiome on transcriptional profiles of the neonatal ileum and the circulating immune compartment. **(B)** Principal component analysis plots demonstrating the distribution of P1 male ileum samples colored by treatment group, revealing clustering is driven by developmental maturity, mode of delivery, and colonization (C-section Amies inoculated vs. vaginally delivered) and the C-section pups inoculated with human cervicovaginal microbiota as an intermediate between the Amies and vaginally delivered pups. P1 corresponds to 24 hrs post-inoculation. N = 3 – 4 males per treatment. **(C)** Heatmap depicting mean expression of genes in the ileum P1 males, showing differences between CST I and CST IV males (linear fit model, FDR < 0.1, log(FC) = 1.5). Unbiased hierarchical clustering showing similarity in transcriptional patterns between CST IV and VD males, and CST I and Amies males. P1 corresponds to 24 hrs post-inoculation. *Z*-scores plotted across individuals for each gene. N = 3 – 4 males per treatment. **(D)** Cluster-based functional enrichment analysis of differentially expressed genes in the ileum of P1 males, showing significant enrichment of pathways involved in defense response to bacteria, innate immune activation, chemotaxis, and mucus secretion in the ileum of CST IV relative to CST I inoculated males (FDR < 0.05). Bubble plot size denotes enrichment. N = 4 – 6 males per treatment. **(E)** t-SNE visualization demonstrating CyTOF phenotyping of CD45+ immune cells in whole blood of CST I, CST IV, and vaginally delivered males showing neutrophils as the major immune subset in the circulating immune compartment at P1 (equal sampling across treatment groups, total 10,000 events). P1 corresponds to 24 hrs post-inoculation. N = 3 males per treatment. **(F)** Average frequencies of major leukocytes within whole blood in P1 CST I, CST IV, and VD males. N = 3 males per treatment. **(G-L)** Frequencies of circulating **(G)** neutrophils, **(H)** monocytes, **(I)** MHC II^+^ B cells, **(J)** MHC II^-^ B cells, **(K)** CD8+ T cells, and **(L)** CD4^+^ T cells within whole blood in P1 CST I, CST IV, and VD males. **(L)** Frequency of CD4^+^ T cells were significantly increased in VD males compared with CST I and CST IV males (One-way ANOVA, F_2,6_ = 15.97, *P* = 0.0040; Tukey’s post-hoc CST I vs. VD *P* = 0.0072; CST IV vs. VD *P* = 0.0059). N = 3 males per treatment. ** *P* < 0.01.

### Exposure to high-risk associated human cervicovaginal microbiota leads to lasting sex-specific changes in post-pubertal growth, circulating immunity, and transcriptional profiles in the hypothalamus

Maternal microbiota colonization in early life influences key aspects of growth, metabolism, immunity, and the development of brain regions involved in the central regulation of homeostatic processes (Cryan et al., 2019; Dinan and Cryan, 2017; McDonald and McCoy, 2019; Vuong et al., 2017, 2020). To further determine the translational relevance of this human microbiota-associated mouse model, we examined the lasting effects of early life colonization by CST I and CST IV on these key microbiota-mediated phenotypes including growth, immunity, and central regulation of metabolism. Body weight measurements from postnatal day (P) 21 to P56 showed that the body weight growth curve for CST IV-exposed males was higher than that of CST I and Amies males, but not VD males (Figure 1C). Comparisons of the percentage of body weight gain from P21 to P56 and the final body weights at P56 revealed significantly different weight trajectories between CST IV-exposed males compared with CST I- and Amies-exposed males (Figure 1C, Supporting Figure 3A,B). No significant effects in rump-snout length were observed between the four groups, indicating body weight differences are independent of body length (Supporting Figure 3C). No differences in body weight measurements were observed between the four groups in female offspring (Figure 1C, Supporting Figure 3D-F). Based on the sex-specific effects of colonization by human cervicovaginal microbiota on growth and metabolism, the proceeding experiments focused on male offspring.

**Figure 3.**
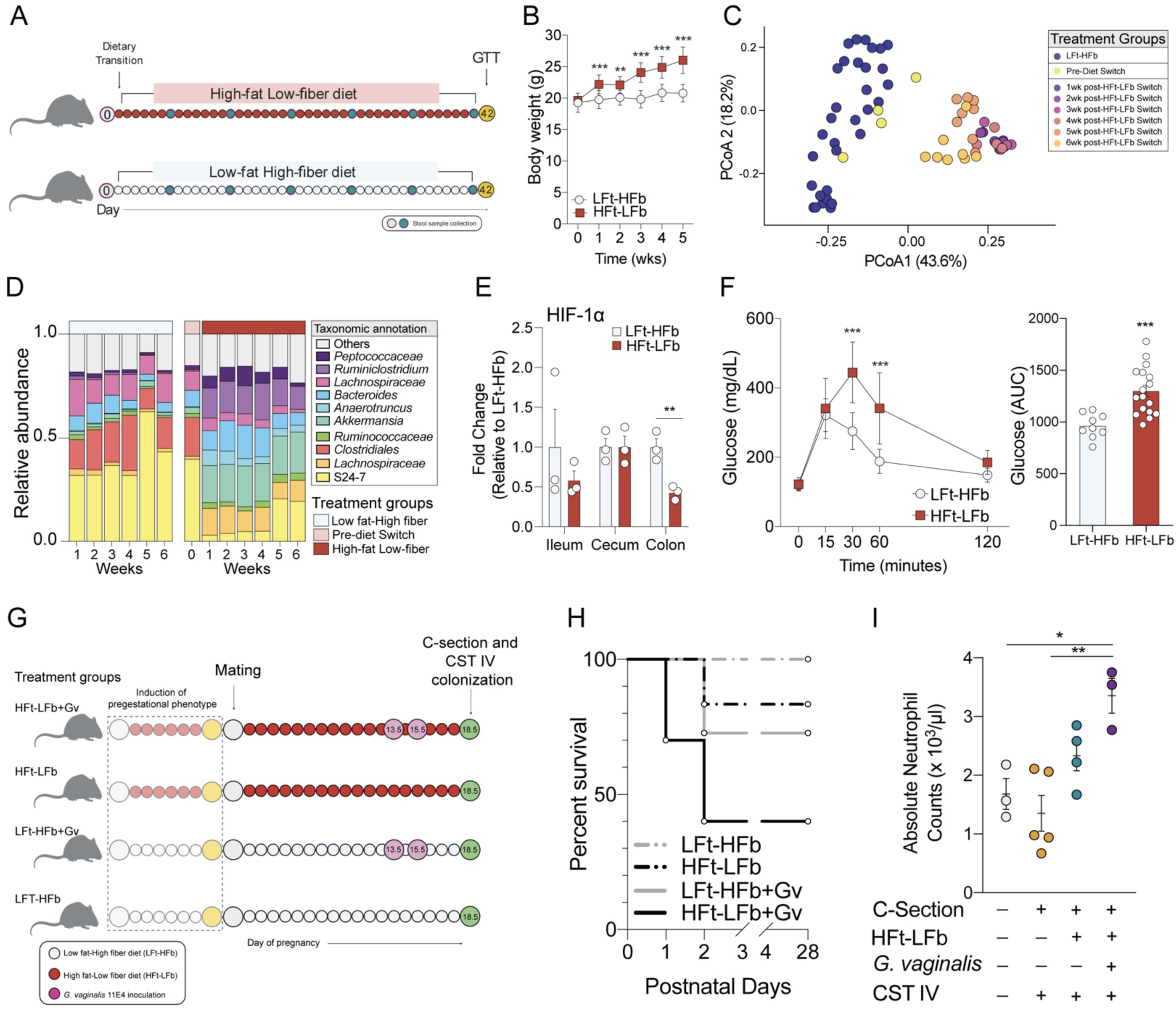
Modeling compounding maternal risk factors increases offspring mortality and an exaggerated offspring innate immune response to human CST IV exposure. **(A)** Schematic of experimental timeline for the induction of pregestational excessive weight gain, glucose intolerance, and microbiota alterations through the consumption of a high-fat low-fiber diet (HFt-LFb). **(B)** Consumption of a high-fat low-fiber diet accelerates body weight gain in females compared with females consuming a low-fat high-fiber diet (two-way ANOVA, main effect of time, F_5, 120_ = 94.17, *P* < 0.0001; main effect of diet, F_1, 24_ = 30.70, *P* < 0.0001; time*diet interaction, F_5, 120_ = 36.61, *P* < 0.0001). N = 12 – 20 females per treatment. Data represented as mean ± SEM. Data is representative of two independent experiments. ** *P <* 0.01, *** *P <* 0.001. **(C)** Principal coordinates analysis demonstrating temporal dynamics of diet on the gut microbiota, whereby 1wk consumption of a high-fat low-fiber diet resulted in separate clustering from females consuming a low-fat high-fiber diet that failed to recover during the treatment window. N = 12 – 20 females per treatment and timepoint, total of 68 samples. **(D)** Mean relative abundance of top ten taxa showing rapid changes to the fecal microbiota following transition to a high-fat low-fat diet, characterized by a loss of Clostridiales. N = 12 – 20 females per treatment and timepoint, total of 68 samples. **(E)** Expression of HIF-1a is significantly decreased in the colon following 6 week consumption of a high-fat low-fiber diet relative to females consuming a low-fat high-fiber diet (*t_4_* = 4.9, *P* = 0.008), indicating possible disruption in hypoxia homeostasis in the colon. ** *P* < 0.01. **(F)** *Left,* Plasma levels of glucose levels during a glucose tolerance test in females consuming either a high-fat low-fiber or low-fat high-fiber diet. Females consuming a high-fat low-fiber diet showed significant delay in glucose clearance (two-way ANOVA, main effect of time, F_4, 108_ = 106.33, *P* < 0.0001; main effect of diet, F_1, 27_ = 15.86, *P* = 0.0005; time*diet interaction, F_4, 108_ = 15.44, *P* < 0.0001). *Right*, AUC of total plasma glucose levels showing increase glucose levels in females consuming a high-fat low-fiber diet (two-tailed T test, *t*_24_ = 4.055, *P =* 0.005). Data represented as mean ± SD with individual datapoints overlaid. *** *P <* 0.001. **(G)** Schematic of experimental design to determine the impact of prenatal exposure to compounding maternal adversities, such as diet and presence of a common member of CST IV, during pregnancy on offspring outcomes. We induced the pregestational phenotype and colonized pregnant females to *G. vaginalis* 11E4 (Gv) gestational day 13.5 and 15.5. At gestational day 18.5, offspring from all treatment groups were colonized with the nonoptimal CST IV human cervicovaginal microbiota. **(H)** Survival of offspring from dams that experience a single or multiple compounding adversities. All pups were C-section delivered and gavaged with human CST IV inoculant, showing the highest offspring mortality risk in HFt-LFb+Gv offspring that were exposed to CST IV at birth. LFt-HFb = low-fat high-fiber; HFt-LFb = high-fat low-fiber; Gv = *G. vaginalis* 11E4. Kaplan-Meier survival analysis. N=20 offspring per treatment condition. **(I)** Compounding effects of maternal diet, maternal vaginal colonization by *G. vaginalis*, and exposure to CST IV on absolute number of neutrophils in the circulation of postnatal day 1 male pups. Triple-hit male pups show significant increase in neutrophils compared with vaginally delivered and CST IV inoculated offspring (One-way ANOVA, main effect of treatment, F_3, 11_ = 8.452, *P* = 0.0034; VD vs. Triple Hit, *P* = 0.0188; CST IV vs. Triple Hit, *P* = 0.0026). N = 3-5 males per group. Data represented as mean ± SEM with individual data points overlaid. *P < 0.05, **P < 0.01

As changes to growth patterns and body weight have been shown to influence the composition and function of the peripheral immune cell populations, we next determined the lasting immunomodulatory effects of CST I and CST IV exposure using single-cell mass cytometry by time of flight (CyTOF) (Rosshart et al., 2019). We assessed changes in the distribution of circulating immune cells in P56 CST I, CST IV, and VD males and observed that exposure to CST I and CST IV was associated with shifts in circulating immune cell composition at P56 (Figure 1D-H; Supporting Figure 4A-C). Specifically, we observed an increase in the frequency of neutrophils in CST I- and CST IV-exposed males and a reduction in B cells (total) and IgD^+^ B cells in CST I males (Figure 1F-H), suggesting the possibility that early-life colonization by distinct constituents of the human cervicovaginal microbiota may exert lasting effects on both the innate and adaptive components of peripheral immunity in a CST-dependent manner.

**Figure 4.**
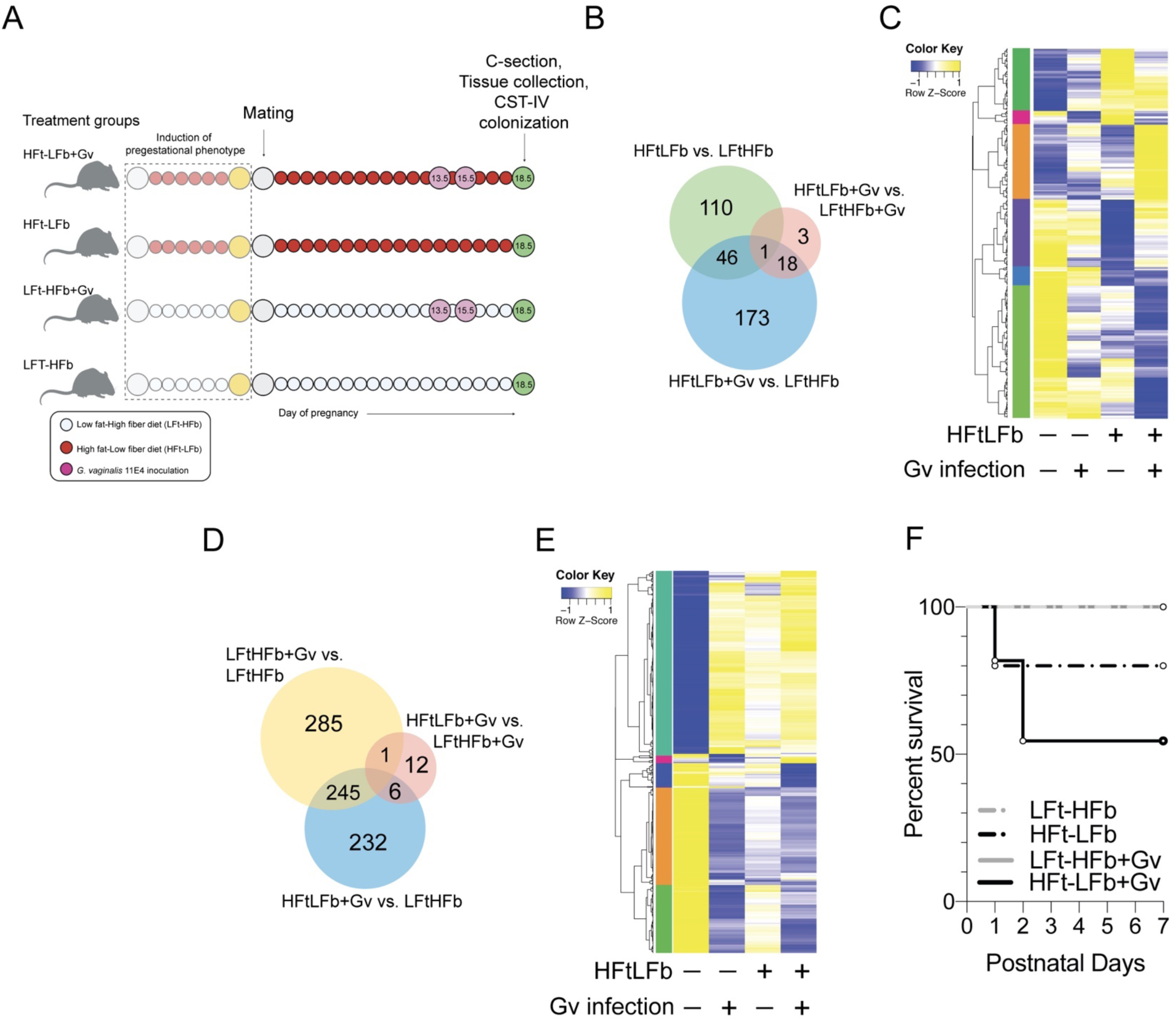
Compounding maternal insults disrupt gene expression patterns in the placenta and fetal ileum that is associated with increased offspring mortality. **(A)** Schematic of experimental design to determine whether compounding maternal adversities, such as diet and *G. vaginalis* vaginal colonization, impact fetal development that may contribute to increased offspring mortality risk. We induced the pregestational phenotype and dams were colonized with *G. vaginalis* 11E4 on gestational day 13.5 and 15.5. At gestational day 18.5, tissue from one cohort of male offspring was collected for analysis of gene expression patterns in the placenta and ileum. A second cohort of offspring were colonized with the nonoptimal CST IV human vaginal microbiota. **(B)** Venn diagram displaying the number of differentially expressed genes in the embryonic day 18.5 placenta of male offspring exposed to a maternal high-fat low fiber diet, *G. vaginalis* 11E4 vaginal colonization, or a combination (linear fit model, FDR < 0.1, log(FC) = 1.5; n = 351 genes). **(C)** Heatmap depicting mean expression of genes in the placenta of embryonic day 18.5 male offspring exposed to a maternal high-fat low fiber diet, *G. vaginalis* 11E4 vaginal colonization, or a combination (HFt-LFb+Gv) (linear fit model, FDR < 0.1, log(FC) = 1.5). These comparisons reveal a unique cluster of genes that are differentially expressed in HFt-LFb+Gv male placenta compared with other treatment groups. *Z*-scores plotted across individuals for each gene. LFt-HFb = low-fat high-fiber; HFt-LFb = high-fat low-fiber; Gv = *G. vaginalis* 11E4. N = 3 males per treatment. **(D)** Venn diagram displaying the number of differentially expressed genes in the embryonic day 18.5 ileum of male offspring exposed to a maternal high-fat low fiber diet, *G. vaginalis* 11E4 vaginal colonization, or a combination (HFt-LFb+Gv) These comparisons reveal a unique cluster of genes that are differentially expressed in the fetal ileum of HFt-LFb+Gv males compared with other treatment groups (linear fit model, FDR < 0.1, log(FC) = 1.5; n = 781 genes). LFt-HFb = low-fat high-fiber; HFt-LFb = high-fat low-fiber; Gv = *G. vaginalis* 11E4. **(E)** Heatmap depicting mean expression of genes in the ileum of embryonic day 18.5 male offspring exposed to a maternal high-fat low fiber diet, *G. vaginalis* 11E4 vaginal colonization, or a combination (linear fit model, FDR < 0.1, log(FC) = 1.5). *Z*-scores plotted across individuals for each gene. N = 3 males per treatment. **(F)** Survival of offspring from dams that experience a single or multiple compounding adversities in a second validation cohort. All pups were C-section delivered and gavaged with human CST IV inoculant, showing the highest offspring mortality risk in HFt-LFb+Gv male offspring exposed to CST IV at birth. LFt-HFb = low-fat high-fiber; HFt-LFb = high-fat low-fiber; Gv = *G. vaginalis* 11E4. Kaplan-Meier survival analysis. N=24-40 offspring per treatment condition.

Given that the hypothalamus is exquisitely sensitive to peripheral shifts in feeding patterns, nutrient availability, metabolism, and immunity, we next compared transcriptional profiles of the paraventricular nucleus of the hypothalamus (PVN) in P56 Amies, CST I, CST IV, and VD males using bulk RNA sequencing (Thaler et al., 2012; Valdearcos et al., 2015; Waterson and Horvath, 2015; Williams et al., 2000, 2001). Differential gene expression analysis (FDR < 0.1, Log(FC) > 1.5) comparing CST I and CST IV males identified 75 genes with altered expression in the PVN (Figure 1I). Unbiased hierarchical clustering of the differentially expressed genes revealed clustering between Amies and CST I males and clustering between CST IV and VD males, paralleling the growth curve and body weight differences observed in these males (Figure 1C,I). Further, cluster-based functional enrichment revealed upregulation of pathways involved in lipid metabolism, insulin receptor signaling, and immune activation in the PVN in CST IV males compared with Amies, CST I, and VD males (Figure 1J). Taken together, these results may suggest that early life colonization by unique constituents of the human cervicovaginal maternal microbiota contribute to lasting metabolic, immune, and neural outcomes in a sex-specific manner. Further, these results point to programmatic effects of colonization by human cervicovaginal maternal microbiota early in life that persist into adulthood.

### Exposure to human cervicovaginal microbiota influences transcriptional programs in the postnatal ileum

Next we sought to determine whether colonization by CST I or CST IV impacts transcriptional profiles of the neonatal ileum at postnatal day 1, given that CST I and CST IV have been shown to induce CST-dependent transcriptional signatures in the female reproductive tract, and that both communities were detected in feces of human infants, (Figure 2A) (Doerflinger et al., 2014). We used bulk RNA sequencing to compare gene expression patterns in the postnatal day P1 ileum of C-section males inoculated by gavage with Amies, CST I, or CST IV as well as age-matched VD male offspring. A principal components analysis showed that Amies and VD males formed distinct clusters, while CST I and CST IV males showed an intermediate phenotype between Amies and VD male offspring (Figure 2B). Differential gene expression analysis (FDR < 0.1, Log(FC) > 1.5) comparing CST I and CST IV males identified 128 genes with altered expression in the P1 ileum (Figure 2C). Unbiased hierarchical clustering of the differentially expressed genes revealed clustering between Amies and CST I males and clustering between CST IV and VD males, suggesting that colonization by CST I or CST IV is associated with unique transcriptional profiles in the P1 ileum (Figure 2C). Cluster-based functional enrichment revealed significant enhancement of metabolic and immune pathways in the ileum of Amies, CST I, CST IV, and VD males (Figure 2D). The CST I enriched cluster included genes enriched in pathways involved in the activation of gasdermin family, necrosis regulation, and stimulation of hormone synthesis. Conversely, the two clusters shared by CST IV and VD males included genes enriched in pathways involved in the regulation of mucus secretion, defense response to bacteria, cholesterol metabolism, neutrophil migration, and chromatin modification (Figure 2D). Our observations that colonization by CST I and CST IV microbiota may induce CST-dependent transcriptional programs in the neonatal mouse ileum is consistent with previous *in vitro* studies showing that colonizing fetal enterocytes with *Lactobacillus* induces a slow and more sustained immune response while colonization by *E*. *coli* induces intestinal tolerance through more rapid immune signaling cascades in fetal enterocytes (Zeuthen et al., 2010). Collectively, these results show that gene expression patterns and functional pathways in the neonate ileum are influenced by the colonizing microbiota in a CST-dependent manner, providing an additional link between early life colonization patterns and our observed lasting outcomes in adulthood.

### Exposure to human cervicovaginal microbiota influences the early life circulating immune compartment

As early life microbiota composition can influence circulating immune cell composition and function, we next determined whether transplantation of CST I or CST IV microbiota influenced circulating immunity in P1 male offspring (Deshmukh et al., 2014; Gensollen et al., 2016; Niu et al., 2020; Olszak et al., 2012; Singer et al., 2019). We used an established model of perinatal antibiotic exposure whereby pregnant dams are exposed to ampicillin, vancomycin, and neomycin to deplete microbiota and thus prevents vertical transmission of maternal microbiota to offspring (Deshmukh et al., 2014). On embryonic day 18.5, offspring were cesarean delivered and male offspring were colonized with CST I, CST IV, or maintained on the antibiotics until P1. Age-matched VD male offspring were also included to account for potential effects of premature birth, mode of delivery, and antibiotic exposure. Hematology analysis revealed that circulating neutrophils were significantly reduced in antibiotic-exposed male offspring compared with CST I, CST IV, and VD male offspring. No differences were observed in circulating neutrophil counts between CST I, CST IV, and VD male offspring (Supporting Figure 5A). Notably, basophil counts were significantly increased in antibiotic-exposed male offspring compared with CST IV and VD male offspring but not CST I offspring, consistent with previous work showing that microbial colonization may regulate circulating basophil populations (Supporting Figure 5B). These results align with previous work showing that colonization by maternal microbiota is necessary for development-dependent expansion and contraction of immune cell populations in the periphery (Deshmukh et al., 2014; Hill et al., 2012; Olin et al., 2018).

A limitation of the hematology analysis is the inability to further classify granulocyte and lymphocyte populations into specific immune subsets. Thus, we determined the immunomodulatory effects of CST I and CST IV microbiota exposure using CyTOF on whole blood from P1 male offspring (Figure 2E-L; Supporting Figure 6 A-C). Given the small volume of whole blood that can be collected from a P1 mouse (∼30 µL per animal), we implemented an established CyTOF workflow using small volumes of whole blood from infants and optimized this procedure for neonatal mice. Consistent with the hematology analysis, this approach revealed that neutrophils are the most abundant immune cell subset in circulation across all groups at postnatal day 1 (Figure 2G). Additionally, monocytes, B cells, macrophages, natural killer cells, CD8^+^ T cells, and CD4^+^ T cells were detected in circulation of P1 CST I, CST IV and VD male offspring (Figure 2H-L). Comparison of immune cell frequencies revealed increased frequency of CD4^+^ T cells in the circulation of VD males compared with CST I and CST IV males, which may reflect developmental differences due to mode of delivery or premature birth in the cesarean delivered and inoculated male offspring (Figure 2L). No other differences in the frequency of circulating immune cell populations were observed. Together, our analysis of the circulating immune compartment revealed multiple immune cell populations that expand and contract in response to early life intestinal microbial colonization, as well as immune cell populations that may be influenced by premature birth and mode of delivery in mice.

### Modeling a high-risk *in utero* environment increased offspring mortality and leads to an exaggerated innate immune response following CST IV colonization at birth

While our mouse model showed that transplanting CST I or CST IV into C-section delivered mouse pups influenced postnatal development, these experiments ignored the possible influence of these cervicovaginal microbiota on prenatal development. In other words, newborn human infants colonized by CST I or CST IV during birth will be exposed to a maternal milieu, and develop within a prenatal environment, that is shaped by the immune and metabolic functions of these microbiota. Previous work from our laboratory has shown that maternal exposures during pregnancy disrupted fetal development, and the consequences of these prenatal exposures persisted across the lifespan (Bronson and Bale, 2014; Howerton et al., 2013; Jašarević et al., 2018; Mueller and Bale, 2008; Nugent et al., 2018). Consistently, recent work supports significant associations between CST IV microbiota, stress, obesity and adverse maternal-child health outcomes.

Given our previously published observations and the literature on how compounding maternal risk factors contribute to offspring adverse outcomes, we next sought to establish a rodent model that may mimic the maternal milieu of women that harbor CST IV and consumed a diet deficient in dietary fibers (Shivakoti et al., 2020). Specifically, we determined whether presence of *G. vaginalis,* a common member of CST IV, and a high fat-low fiber diet impacted offspring health outcomes. In this model, prepubertal females were transitioned to consume a high-fat low-fiber (HFt-LFb) diet or continued to consume a low-fat high-fiber (LFt-HFb) diet at postnatal 28 (Figure 3A). Weekly body weight measurements across a period of six weeks showed a significant body weight increase in HFt-LFb females compared with LFt-HFb females (Figure 3B). Body weight changes accompanied significant shifts in gut microbiota structure and composition, characterized by an enduring loss of the soluble fiber fermenting members of the class *Clostridia* and a bloom of *Akkermansia* in HFt-LFb relative to LFt-HFb females (Figure 3C,D). Differential abundance analysis revealed disruption in the abundance of 48 taxa between HFt-LFb and LFt-HFb females (**Supporting Table 2**). Consistent with previous work showing loss of intestinal hypoxia due to high-fat low-fiber diet consumption, we found decreased expression of the hypoxia-inducible factor 1-alpha in the colon of HFt-LFb relative to LFt-HFb females (Figure 3E) (Byndloss et al., 2018; Faber and Bäumler, 2014; Lee et al., 2020). Further, consumption of a high fat-low fiber diet has been associated with disruption to glucose homeostasis in females (Gohir et al., 2019; Montgomery et al., 2013). At seven weeks of diet consumption, glucose clearance was significantly delayed in HFt-LFb relative to LFt-HFb females following administration of a glucose tolerance test (Figure 3F).

We next investigated the effects of HFt-LFb consumption on maternal outcomes during pregnancy. Excessive body weight gain was observed in dams consuming a HFt-LFb from gestational day 0.5 to 14.5 compared with LFt-HFb dams. Differences in body weight disappeared between gestational day 15.5 to 18.5, which was attributed to a change in the trajectory of daily body weight gain in HFt-LFb dams relative to LFt-HFb dams (Supporting Figure 7A,B). Consistent with disruption to maternal body weight trajectories, daily tracking of the maternal gut microbiota revealed significant alterations to microbiota structure and composition in HFt-LFb dams compared with LFt-HFb dams across pregnancy (Supporting Figure 7C,D). Differential abundance analysis comparing the relative abundance of gut microbial taxa across pregnancy between HFt-LFb and LFt-HFb revealed 36 differentially abundant taxa. Of these differentially abundant taxa, the relative abundance of butyrate-producing *Butyricoccus* and the immunomodulatory *Candidatus* Arthromitus was reduced in HFt-LFb compared with LFt-HFb pregnant dams (Supporting Figure 7E,F) (Geirnaert et al., 2014; Ivanov et al., 2009).

After establishing the effect of diet on maternal body weight, glucose homeostasis, and gut microbiota, pregnant HFt-LFb or LFt-HFb dams were intravaginally inoculated with 2×10^8^ colony forming units of *G. vaginalis* strain 11E4 (Amar et al., 2007; Andersen et al., 2016; Gilbert et al., 2013; Milner and Beck, 2012; Sierra et al., 2018). This bacterial species was chosen based on previous work showing that *G. vaginalis* is a common member of CST IV, induces immune activation and inflammation in the cervicovaginal space and contributes to the pathogenesis of bacterial vaginosis in humans and rodent models (Gilbert et al., 2013; Romero et al., 2004; Sierra et al., 2018). Inoculating HFt-LFb or LFt-HFb dams with *G. vaginalis* 11E4 at gestational day 13.5 and 15.5 did not impact body weight or gut microbiota (Figure 3A-F; Supporting Figure 7). As further validation, we confirmed presence of *G. vaginalis* in the vaginal fluid of inoculated compared with non-inoculated dams (Supporting Figure 8A).

Given that prenatal exposures have been shown to impact the ways in which offspring respond to the postnatal environment, we next determined whether maternal HFt-LFb and *G. vaginalis* vaginal colonization, alone or in combination, influenced offspring response to being exposed to CST IV at birth. C-section delivered LFt-HFb, HFt-LFb, LFt-HFb+Gv, and HFt-LFb+Gv offspring were exposed to CST IV, transferred to foster dams, and monitored for litter acceptance and survival (Figure 3G). Following colonization with CST IV, HFt-LFb+Gv male offspring showed a 60% mortality rate by postnatal day 3 compared with the 29% mortality in LFt-HFb+Gv males, 17% in HFt-LFb males, and no mortality in LFt-HFb (Figure 3H). No additional mortality was observed beyond postnatal day 3 (Figure 3H). To determine whether increased offspring mortality risk was specific to CST IV, we also transplanted CST I into a separate cohort of offspring with exposed to the varying combinations of maternal diet and *G. vaginalis* vaginal colonization. We observed that HFt-LFb+Gv male offspring colonized with CST IV exhibited a mortality rate of 60% while HFt-LFb+Gv male offspring colonized with CST I exhibited a mortality rate of 50%, showing an 10% increase in the survival rate among high-risk offspring colonized by CST I in this mouse model (Supporting Figure 9).

Given our observations that early life transplantation of human cervicovaginal microbiota is associated with increased circulating neutrophils, we next determined whether increased mortality risk in HFt-LFb+Gv male offspring is related to neutrophil counts at postnatal day 1. Similar to our previous experiments, age-matched VD offspring were included as a baseline comparison for mode of delivery, premature birth, and colonization in C-section delivered offspring. Neutrophil counts were significantly increased in HFt-LFb+Gv male offspring colonized with CST IV compared to LFt-HFb males colonized with CST IV and VD males, while no difference was detected between HFt-LFb and HFt-LFb+Gv male offspring colonized with CST IV (Figure 3I). Specifically, neutrophil counts more than doubled between HFt-LFb+Gv (Mean = 3.5 x 10^3^ per µL of blood) and LFt-HFb (Mean = 1.5 x 10^3^ per µL of blood), consistent with a clinical definition of neutrophilia in HFt-LFb+Gv male offspring. Overall, these results suggest that prenatal factors have a profound effect on determining offspring response to the postnatal environment, including the immune response to exposure to maternal cervicovaginal microbiota.

### Compounding maternal risk factors disrupt gene expression patterns in the placenta and fetal ileum that is associated with increased offspring mortality risk

Given our observations regarding compounding maternal risk factors and increased offspring mortality risk, we next determined whether prenatal exposure to a maternal HFt-LFb diet and *G. vaginalis* vaginal colonization disrupted development of the fetal compartment. An independent cohort of pubertal females were transitioned onto the high fat-low fiber diet, maintained on the assigned diets through pregnancy, and infected with *G. vaginalis* 11E4 during pregnancy. To validate the reproducibility of the pregestational phenotype, we replicated the excessive weight gain, delayed glucose clearance, and increased circulating glucose levels in females following consumption of a high fat-low fiber for six weeks (Supporting Figure 10). Our analysis was focused on the placenta and ileum based on previous reports showing that the placenta and fetal ileum are particularly sensitive to the disruptive effects of maternal diet and nonoptimal cervicovaginal microbiota (Gohir et al., 2019; Kemp, 2014). Gene expression patterns of the placenta and ileum were analyzed in embryonic day 18.5 male offspring from LFt-HFb dams, male offspring from LFt-HFb dams infected with *G. vaginalis* (LFt-HFb+Gv), male offspring from HFt-LFb dams (HFt-LFb), and male offspring from HFt-LFb dams infected with *G. vaginalis* (HFt-LFb+Gv) by bulk RNA sequencing (Figure 4A).

In the placenta, differential gene expression analysis (FDR < 0.01, log(FC) = 1.5) revealed that nearly one-third, 110 of 351, of genes with significant differences were altered in the HFt-LFb male placenta compared with LFt-HFb male placenta, with the majority of these genes downregulated in HFt-LFb relative to LFt-HFb male placentas (Figure 4B,C). Functional enrichment of these HFt-LFb-specific downregulated genes enrich in pathways involved in the response to changing nutrient levels, fatty acid metabolism, dopamine transport, and serotonin transport. Additionally, 173 of 351 genes were differentially expressed between HFt-LFb+Gv and LFt-HFb male placenta. Functional enrichment of these differentially expressed genes show enrichment in pathways involved in collagen degradation, response to inflammation, and receptor-mediated endocytosis. Presence of *G. vaginalis* in the maternal cervicovaginal space was associated with a cluster of differentially expressed genes in the male placentas in a diet-independent manner. These genes were enriched in pathways involved in glucose homeostasis and fatty acid oxidation (Supporting Figure 11A). No gene expression differences between LFt-HFb+Gv and LFt-HFb male placentas passed FDR cut-off.

Moreover, gene set enrichment analysis (GSEA) was used to identify pathways that are altered by *G. vaginalis* 11E4 vaginal colonization, a high fat-low fiber diet, or in combination. Comparison of gene sets showed significant enrichment of pathways involved in the activation of innate immune responses, interleukin-6 biosynthetic processes, and negative regulation of interleukin-10 production in the LFt-HFb+Gv placenta compared to male placenta from LFt-HFb dams. These results show that presence of *G. vaginalis* in the maternal cervicovaginal space has significant effects on gene expression patterns in the E18.5 male placenta (Supporting Figure 12A). Male placenta samples from HFt-LFb dams showed significant disruption in pathways involved metabolic processes, including steroid metabolic processes and oxidative phosphorylation (Supporting Figure 12B). Further, comparison between HFt-LFb+Gv and HFt-LFb male placentas revealed significant disruption in adaptive immune response pathways in HFt-LFb+Gv placentas, pointing to a transcriptional signature of an impaired immune response in the HFt-LFb male placenta following maternal vaginal exposure to *G. vaginalis* (Supporting Figure 12C). Consistent with this notion of impaired maturational processes, comparisons between HFt-LFb+Gv and LFt-HFb males revealed significant disruption to pathways involved in the regulation of circulatory processes and regulation of vasculature development in the placenta of HFt-LFb+Gv males relative to that of LFt-HFb males (Supporting Figure 12D).

As alterations to placental handling of nutrients, metabolites, and oxygen transport may influence the development of fetal tissues, we used bulk RNA sequencing to determine the additive effect of maternal diet and *G. vaginalis* vaginal colonization on the fetal ileum. Differential gene expression analysis (FDR < 0.1, log(FC) = 1.5) revealed that 285 of 781 differentially expressed in the fetal ileum of LFt-HFb and LFt-HFb+Gv males (Figure 4D,E). Additionally, 232 of 781 genes were differentially expressed between HFt-LFb+Gv and LFt-HFb male ileum (Figure 4D,E). We also found a cluster of genes that were downregulated in the ileum of HFt-LFb+Gv males relative to LFt-HFb, LFt-HFb+Gv, and HFt-LFb males. These downregulated genes were enriched in pathways involved in oxygen transport, antigen processing, cholesterol metabolic processes, lymph node development, heme metabolic process, and regulation leukocyte migration (Supporting Figure 11B), suggesting disruption of important maturational processes in the ileum that only occurs in the combinatorial exposure to maternal HFt-LFb and *G. vaginalis* vaginal colonization. No gene expression differences between male placentas across treatment groups passed FDR cut-off.

Paralleling the broad disruption to metabolic and immune pathways in the placenta, GSEA showed significant alterations in the fetal ileum due to maternal diet, infection, and their combination. Comparison of gene sets showed significant enrichment of pathways involved in immune activation and acute inflammatory responses in the LFt-HFb+Gv male fetal ileum compared with LFt-HFb male ileum (Supporting Figure 13A). Male ileum samples from HFt-LFb dams showed significant disruption in maturational pathways, including epithelial cell development (Supporting Figure 13B). Further, comparison between HFt-LFb+Gv and HFt-LFb male ileum revealed significant disruption in the regulation of vasculature development, epithelial cell proliferation, and myeloid mediated immunity in HFt-LFb+Gv ileum, consistent with disruption to critical maturation of tissue processes (Supporting Figure 13C). Collectively, these results showed that presence of vaginal *G. vaginalis* and maternal consumption of a HFt-LFb diet during pregnancy resulted in broad transcriptional dysregulation in the placenta and ileum that was not observed in either treatment alone. This transcriptional signature in the HFt-LFb+Gv placenta and ileum reflect impairment of pathways that support critical maturational and developmental processes and revealed further programmatic effects that may be the direct consequence of compounding maternal risk factors. Lastly, to determine whether these broad disruptions to the gene expression patterns in the placenta and ileum were associated with offspring mortality risk, a second cohort of embryonic day 18.5 LFt-HFb, HFt-LFb, LFt-HFb+Gv, and HFt-LFb+Gv offspring were cesarean delivered and colonized with the nonoptimal CST IV human cervicovaginal microbiota. Replicating the results from our discovery cohort, HFt-LFb+Gv offspring colonized with CST IV showed the highest mortality rate when compared with LFt-HFb, HFt-LFb, LFt-HFb+Gv male offspring colonized with CST IV (Figure 4F).

## Discussion

A nonoptimal CST IV cervicovaginal microbiome is common in Black women and has been associated with negative maternal-child health outcomes, including obstetric complications, spontaneous preterm birth, and neonatal morbidity and mortality (Brown et al., 2018, 2018; Callahan et al., 2017; DiGiulio et al., 2015; Elovitz et al., 2019; Fettweis et al., 2019; Florova et al., 2021; Gerson et al., 2020a; Hočevar et al., 2019; Ravel et al., 2011; Romero et al., 2014; Tabatabaei et al., 2019, 2019; Zhou et al., 2007). This underlying biological risk is further amplified by a variety of environmental factors that contribute to a high risk *in utero* environment, including, but not limited to, obesity, diabetes and consumption of a diet that deficient in dietary fibers (Brookheart et al., 2019; Danese and Lewis, 2017; Gerson et al., 2020b; Shivakoti et al., 2020). While these maternal environmental perturbations alter the composition of the microbiota transferred from mother to offspring, the possibility that prenatal exposure to these risk factors may impact developmental processes that support offspring response and adaptation to postnatal colonization remains unclear.

To address these complex interactions, we first determined whether vertical transmission of human cervicovaginal microbiota contributed to lasting offspring outcomes in a mouse model. We used our previously-established mouse model in which pups were cesarean delivered at embryonic day 18.5 to prevent vaginal colonization and then inoculated by oral gavage with human CST I or CST IV cervicovaginal samples obtained from pregnant women (Jašarević et al., 2018). We observed transient colonization of the human cervicovaginal microbiota in the mouse intestinal tract through confirmation of successful inoculation. As offspring from all groups shared the same postnatal environment, this approach permitted examination as to how early life intestinal colonization by the human cervicovaginal microbiota had lasting effects on key aspects of growth, immunity, and brain development. Colonization by CST I and CST IV showed a sex-specific effect on body weight trajectories, where females showed no differences in body weight, but CST IV inoculated males showed a significant increase in body weight. We further observed shifts in circulating immune composition and changes to gene expression patterns in the paraventricular nucleus of the hypothalamus in CST IV adult males, consistent with altered feeding patterns and metabolic regulation that manifests in changes to growth trajectories and body weight.

These lasting CST-dependent effects of the maternal cervicovaginal microbiota on offspring phenotypic outcomes pointed to possible programmatic effects of these microbiota in early life. A fundamental challenge for the newborn while transitioning from prenatal to postnatal environments is the necessity to balance mounting an appropriate defense response, the energetic costs required to mount a response, and minimizing the level of damage that may stem from that immune response (Harbeson et al., 2018; Kollmann et al., 2017; Macpherson et al., 2017; Renz et al., 2018). As one of the primary sites of microbial colonization, the neonate intestinal tract has a number of transcriptional, metabolic, and immune processes that induce rapid tolerance to prevent an exaggerated inflammatory response during early life colonization (Bittinger et al., 2020; Harbeson et al., 2018; Kollmann et al., 2017; Renz et al., 2018). Activation of these processes is delayed in cesarean delivered mice and missing this window of opportunity is associated with increased susceptibility to infection across the lifespan (Lotz et al., 2006; Morais et al., 2020). Indeed, bulk RNA-seq analysis on postnatal day 1 ileum from Amies, CST I, CST IV, and VD males revealed gene expression patterns and pathways that were modulated by mode of delivery, developmental maturity, and cervicovaginal CST. The postnatal ileum of cesarean delivered males inoculated with CST IV showed significant enrichment of pathways involved in mucus secretion, defense response, and leukocyte recruitment compared with CST I males. Presence of CST IV and CST I has been shown to stimulate CST-dependent transcriptional and immune effects in the female reproductive tract, suggesting the possibility that colonization by CST I and CST IV induces similar pathways in the postnatal ileum in mouse offspring (Anahtar et al., 2015; Doerflinger et al., 2014; Gosmann et al., 2017). As vaginally delivered mice are colonized by *Gammaproteobacteria* have been shown to induce a potent immune response in newborn mice, these community-dependent differences in the induction of an offspring response to colonization may provide one possible explanation for our observed similarities between CST IV and vaginally delivered male offspring. Nevertheless, this model now permits further assessment to identify the specific microbiota-derived signals from CST I and CST IV involved in the induction of these transcriptional programs (Jašarević et al., 2017; Mirpuri et al., 2014).

Recent advances in high-dimensional, single-cell systems-level analyses have made it possible to monitor how early life environmental exposures shape immune system development (Olin et al., 2018). Single-cell mass cytometry optimized for small volumes of whole blood from CST I, CST IV, and VD males revealed that neutrophils encompass the largest component of the circulating immune compartment at postnatal day 1. Expansion of circulating neutrophils occurred in parallel with contraction of circulating basophils. This pattern is consistent with previous work that has also demonstrated the important role of microbial colonization in regulating neutrophil and basophil counts in circulation (Deshmukh et al., 2014; Hill et al., 2012). Differences in the frequency of CD4^+^ T cells between vaginally delivered males and cesarean delivered males may highlight important contributions of mode of delivery and developmental maturity to circulating T cell populations (Olin et al., 2018). Importantly, as the human cervicovaginal microbiota transiently colonizes the infant gut and is replaced by more stably colonizing microbiota from the maternal gut, additional studies are now possible to examine the causes and consequences of subsequent colonization by maternal human skin, gut, and breastmilk-associated microbiota in this model (Ferretti et al., 2018; Yassour et al., 2018). The methods reported here provide additional avenues to examine the mechanistic role of maternal microbiota on immune system maturation, metabolic processes, brain development, and how these early life events may predispose to disease later in life.

To determine whether a high-risk prenatal environment modulate the offspring response to colonization by maternal cervicovaginal microbiota, we established a mouse model of compounding maternal exposures in which offspring are exposed *in utero* to maternal high-fat low-fiber diet and *G. vaginalis* vaginal colonization. Using this model, we revealed that consumption of a high-fat low-fiber diet altered maternal body weight, glucose tolerance, and gut microbiota composition. Consistent with previous studies, consumption of a high fat-low fiber diet reduced relative abundance of gut microbes involved in fermentation, production of short-chain fatty acids, and microbes that exhibit potent immunomodulatory capacity (Gohir et al., 2019). We determined the impact of maternal consumption of a high fat-low fiber diet on fetal development, either in the presence or absence of maternal *G. vaginalis* vaginal colonization. We observed broad disruption to gene expression patterns in the placenta and fetal ileum and identified transcriptional profiles that were specific to offspring from dams consuming a high fat-low fiber diet and colonized by *G. vaginalis*. These offspring showed significant disruption gene network involved in maturational and developmental processes, revealing novel pathways that appear only when modeling multiple maternal risk factors in parallel. One consequence of this developmental immaturity is an excessive influx of neutrophils following intestinal colonization by the nonoptimal CST IV at birth. Further, this heightened innate immune response was accompanied by a significant increase in mortality risk in these offspring. Importantly, we observed increased survival when offspring were colonized with CST I, suggesting that exposure to the nonoptimal CST IV cervicovaginal microbiome may exacerbate negative outcomes in high-risk premature offspring.

Additional work will be needed to identify the mechanisms by which prenatal exposure and maternal microbiota interact to confer susceptibility to offspring morbidity and mortality, as well as lasting outcomes across the lifespan. Nevertheless, should this mouse model be translationally relevant for humans, our findings highlight that pervasive and compounding maternal exposures, such as poor diet and nonoptimal cervicovaginal microbiota, may impair the development of fetal tissues that function as first-responders to postnatal microbial colonization. Our results also have implications for better estimating the combined risk of biological and environmental factors and the ways in which these health disparities contribute to offspring morbidity and mortality. Given the modifiable nature of the maternal microbiota composition and function through possible dietary interventions, our studies may point to unique prenatal and postnatal periods for intervention that may yield novel therapies and biomarkers to be further examined in clinical trials.

## Acknowledgments

The authors thank Mike Humphrys from the Institute for Genome Sciences, Microbiome Service Laboratory at the University of Maryland School of Medicine for assistance with 16S rRNA gene sequence profiling. We thank Kevin Brown from Fluidigm for assistance with development of the CyTOF antibody panel, Michael Solga and Claude Chew from the University of Virginia School of Medicine with acquisition of single-cell mass cytometry data, and El-ad David Amir and the Astrolabe platform for technical assistance with CyTOF data analysis. We also thank Hannah Zierden and Yasmine Cisse for insightful discussion and critical feedback on the manuscript. Research reported in this publication was supported in part by *National Institute of Mental Health, Eunice Kennedy Shriver National Institute of Child Health and Human Development, National Institute of Environmental Sciences, National Institute of Environmental Health Sciences, National Institute of Diabetes and Digestive and Kidney Diseases, and the Institute for Nursing Research* of the National Institutes of Health under award numbers R01MH073030 (TLB), R37MH108286 (TLB), R01HD097093 (TLB), R01ES028202 (TLB), K99HD091376 (KEM), R01NR014826 (JR), R01NR015495 (JR), F32MH109298 (EJ) and K01DK121734 (EJ). The content is solely the responsibility of the authors and does not necessarily represent the official views of the National Institutes of Health.

## Author contributions

Conceptualization, E.J., J.R., and T.L.B; Methodology, E.J., J.R., and T.L.B. Formal Analysis, E.J., K.E.M., T.G., E.H., K.R.; Investigation, E.J., P.J.K., E.H., C.D.H., L.R., K.E.M., L.F., T.G.; Resources, E.J., J.R., and T.L.B.; Writing, E.J., T.L.B; Visualization, E.J.; Supervision, T.L.B; Project Administration, E.J. and T.L.B.; Funding Acquisition, E.J. and T.L.B.

## Declaration of interests

Jacques Ravel is Founder and Chief Scientist at LUCA Biologics. The remaining authors declare no competing interests.

## KEY RESOURCES TABLE

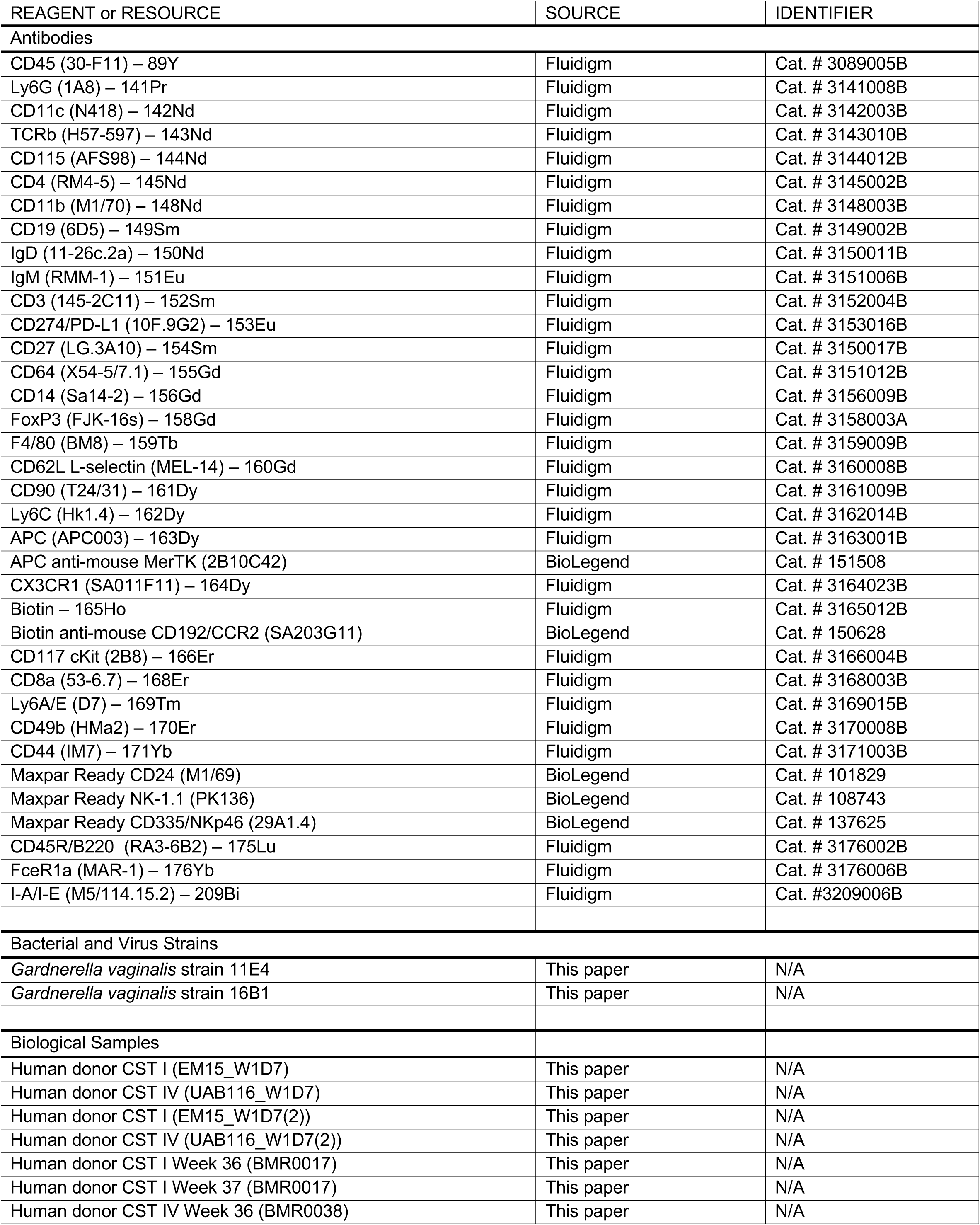
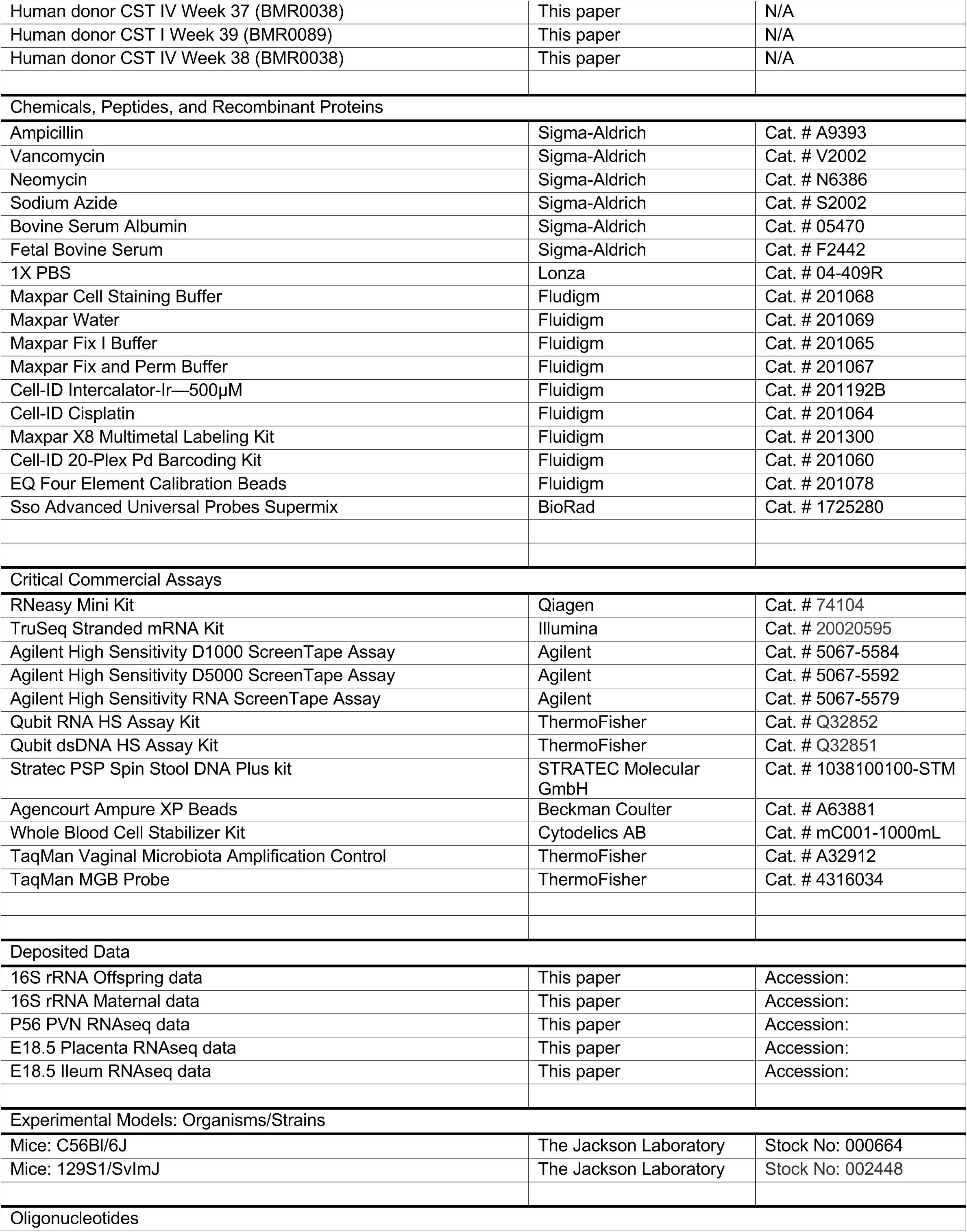
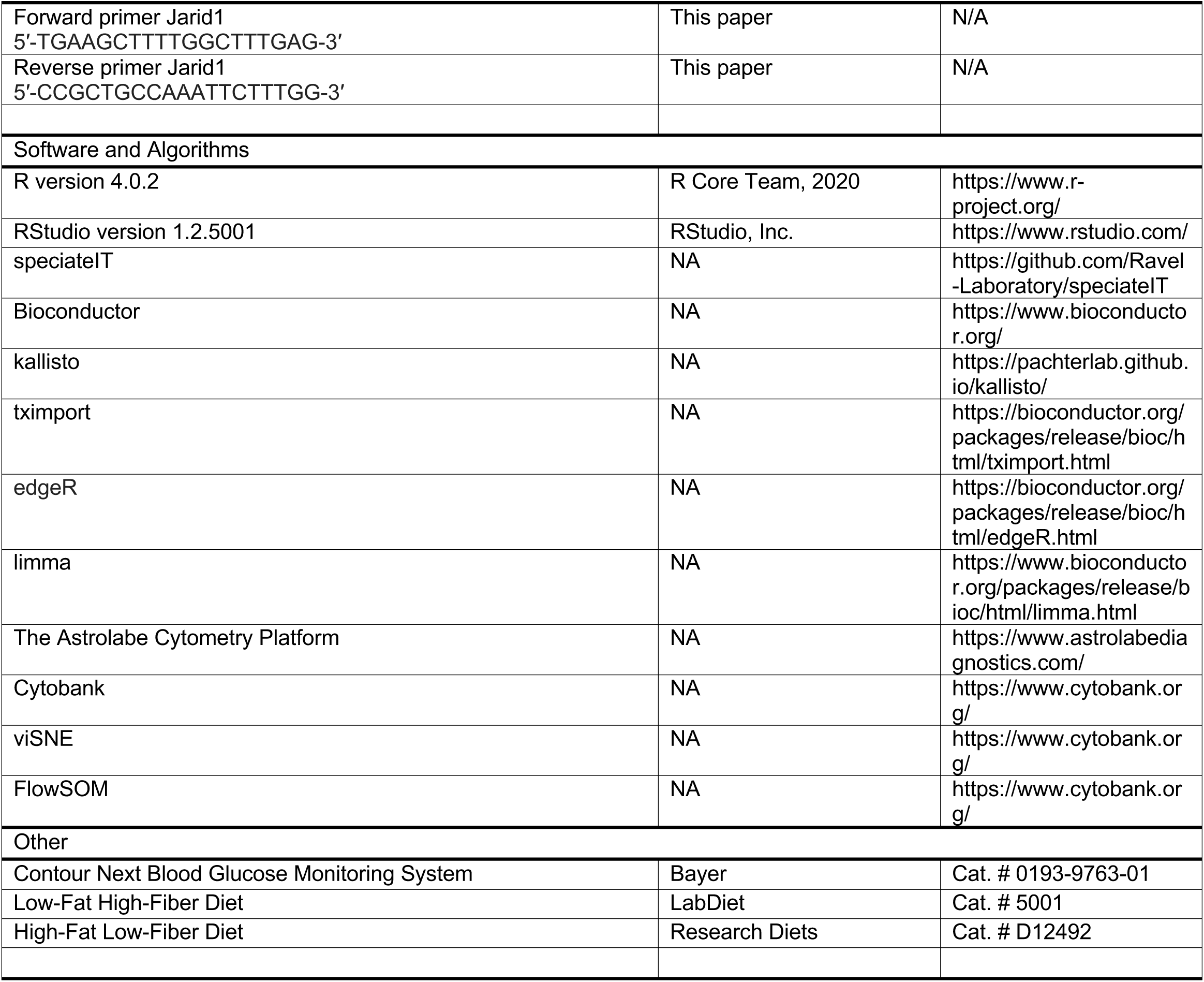

## Experimental Model Details

### Human samples

Human vaginal secretions were selected from a collection of vaginal swabs self-collected weekly until delivery by pregnant women enrolled in the Birth, Eating, and the Microbiome (BEAM) study at the University of Maryland Baltimore. The clinical study protocol (HP-00056389) was approved by the Institutional Review Board of the University of Maryland Baltimore. Written informed consent was appropriately obtained from all participants.

Samples were self-collected using the Copan ESwab system and stored frozen at −20°C in 1ml of Amies Transport Medium to preserve cervicovaginal microbiota composition until transport to the laboratory where the samples were stored at −80°C. As part of the parent study the vaginal microbiota composition of all samples collected was established using previously validated protocols (Holm et al., 2019). CST were established using VALENCIA, a nearest centroid classification method for vaginal microbial communities based on composition (France et al., 2020). Sample selection criteria for transplantation experiments included: 1) harboring a CST I or CST IV vaginal microbiota, 2) availability of samples between 36-39 weeks of pregnancy, and 3) women remained in the respective CST throughout pregnancy. The microbiota-seeded in Amies transport medium was used as the inoculant (20µL).

### Mice

Mice were housed under a 12 h light/day photoperiod with lights on an 0700 EST and ad libitum access to water and a grain-based chow diet (Purina Rodent Chow, St. Louis, MO; 28.1% protein, 59.8% carbohydrate, 12.1% fat). Additional details on diet manipulation experiments appear below. Copulation plugs were checked within 2-hours of “lights on” in order to most accurately estimate the number of animals that were going to be used in the study. Noon on the day that the plug was observed was considered embryonic day 0.5 (E0.5). Samples from one male and one female per litter were used for all subsequent analyses. All experiments were approved by the University of Maryland School of Medicine Institutional Animal Care and Use Committee and performed in accordance with National Institutes of Health Animal Care and Use Guidelines.

### Cesarean delivery surgery

C-section surgeries were conducted as previously described (Jasarevic et al., NN). Donor mice used in these studies were C57Bl/6J females ordered from JAX laboratories at 4 weeks old. Foster mice used in these studies were on a C57BL/6:129 background that are maintained in-house as an outbred mixed colony. This mixed colony is used as the foster based on well-documented occurrence of infanticide of non-genetically related offspring in C57Bl/6J dams. In published and current studies using mixed colony females has yielded a 100% acceptance rate of foster animals (Jašarević et al., 2018). Foster females were time-mated 24h before the expected surgery of donor females (E18.5). Foster females delivered a day prior to donor to ensure successful acceptance of the fostered litter.

Cesarean deliveries of offspring were conducted under sterile conditions, and environmental swabs were taken from surgery suite area to account for contamination. For the surgery, immediately following rapid decapitation, dams were submerged in 70% EtOH for 30 s to minimize contamination of the uterine horn from maternal skin bacteria during extraction. Following removal, uterine horns were placed on a sterile field inside a sterile incubator. Once the fetus and placenta were excised from the uterine horn, the placenta and membranes that surround pups were removed and the umbilical cord was ligated. The remaining membranes and fluid from nose and mouth of the pup were cleared, and after breathing was stimulated, pups were kept warm on a sterile field inside a separate sterile incubator (Rodent Warmer x1, Braintree Scientific Inc., MA).

Offspring with overt signs of developmental delay, defined as small body size relative to litters or presence of a neighboring resorption site, were excluded from experiments. Time from maternal euthanasia to stimulation of offspring breathing was 2 minutes across all experiments. All pups were checked for viability prior to microbial transplantation by observance of healthy skin color and toe pinch withdrawal reflex. Time from stimulation of breathing to transplantation of human microbiota was 10 minutes.

### Mouse Colonization

On the day of colonization experiments, 20 µL of selected human vaginal secretion samples stored in Amies transport medium were retrieved from the −80°C and thawed on ice. Upon thawing, samples were briefly centrifuged to ensure any residual liquid was pelleted. Samples remained on ice during surgeries. Orogastric gavage occurred by guiding a silastic catheter (Intramedic Polyethylene Tubing PE 10, Becton Dickinson, Franklin Lakes, NJ) into the neonate’s esophagus, upon which offspring were fed once with 20 µL of fluid containing human cervicovaginal microbiota. Pup vital signs were checked at this time. All animals were checked daily following surgery for the presence for a milk dot, indicating successful feeding and acceptance of pup by the foster mother.

For experiments designed to examine the effect of postnatal colonization by human cervicovaginal microbiota on offspring outcomes, C-section pups were randomly assigned to be inoculated with Amies transport medium only (Amies), CST I or CST IV groups. Vaginally delivered offspring were born within 0.5 days as C-section delivered pups. Given the differences in developmental maturity between vaginally delivered pups relative to C-section pups, comparisons between vaginal delivered and C-section pups were only conducted for validation experiments in which we examined lasting consequences of premature birth and colonization with human cervicovaginal microbiota on outcomes in mice. While donor females were treatment naïve, we added an additional control for maternal effects by using at least four donor dams as the pool for offspring for each round of surgeries. Surgeries were repeated for a total of five times for early life and adult timepoints. For statistical analysis, the individual offspring constituted the data point in these experiments, and this is reflected in figure legends. For experiments designed to examine the effect of compounding maternal insults and postnatal colonization by human cervicovaginal microbial communities on offspring outcomes, the order of C-section delivery was randomized for LFt-HFb, LFt-HFb+Gv, HFt-LFb, and HFt-LFb+Gv dams, thus yielding all groups for each round of surgery. Surgeries were repeated across four independent biological replicates. For statistical analysis, the litter constituted the data point in these experiments, and this is reflected in figure legends. Finally, to control for the well-documented within-cage effects of postnatal environment on measured outcomes, offspring from all treatment groups were tattooed (Black animal tattoo paste, Ketchum Mfg. Co.) to identify treatment group and then fostered to the same foster dam. One experimenter conducted the tattooing while two separate experimenters that were blind to group treatments conducted all of the postnatal assessments.

### Assessments in C-section Offspring

For offspring, weekly body weights were collected starting at postnatal day 7 and fecal pellets were collected starting at postnatal day 21 to examine the impact early life colonization on growth and microbiota maturation. On postnatal day 56, male and female adults were sacrificed, and body measurements, intestinal segments, blood, and brain samples were collected. Tissues were processed as described below.

## Method Details

### Early life microbiota source tracking

To examine the possible origins of microbiota present in the postnatal intestinal tract of C-section delivered offspring, we used a source tracking approach in which we collected samples from the local cage environment and the foster dam. To account for microbial density in the cage environment, foster dam cages were not changed within one week of C-section cross-fostering experiments. Local cage environment samples included corn cob that surrounded the maternal nest, and the cotton bedding with which the pup litter had direct contact. Dried fecal pellets were removed from both environmental samples to ensure capturing sample-specific microbial community composition. Maternal samples included maternal oral, skin swabs of the nipple chain, cecal, colon, and vaginal. Offspring samples included milk dot, ileum, jejunum, ileum and cecum, and colon samples. All samples were processed as described below.

### Diet-induced pregestational body weight gain and glucose intolerance

An important note on treatment group notation: For these experiments we explicitly highlight the low soluble fiber in commercially available refined high fat diet formulations due to accumulating evidence that the lack of soluble fiber in these dietary formulations is an important contributor to excessive weight gain, obesity, and diabetes in mouse models of diet-induced metabolic syndrome and obesity (Dalby et al., 2017; Morrison et al., 2020). Thus, following a two-week acclimation period consuming the Low Fat – High Fiber (LFt-HFb) diet, six week old C57Bl/6J females were randomly assigned to either transition on the High Fat – Low Fiber (HFt-LFb; ResearchDiets Inc, D12492) diet or remain on the Low Fat – High Fiber (LFt-HFb) diet (grain based chow Purina 5001). Females had *ad libitum* access to respective diets for the span of the experiment. To confirm a pre-diabetic state in HFt-LFb females, glucose tolerance test was administered in 12 week old C57Bl/6J females following six weeks on either HFt-LFb or LFt-HFb diets. Food was removed at 0800 EST and mice were fasted for 6 hours, upon which mice were injected intraperitoneally with 0.3g/mL glucose in saline. Glucose readings were collected by tail blood at 0, 30, 60 and 120 minute timepoints using the Contour Next Blood Glucose Monitoring System (Bayer Co, Germany). Pilot experiments revealed that ∼15% of females consuming the HFt-LFb diet were resistant to diet-induced body weight gain and intolerance. As a result, we established the following inclusion criteria: females on the respective diets do not overlap by 2 standard deviations on body weight and glucose tolerance test were excluded from experiment before breeding and subsequent treatments. Females remained on the assigned diets for the span of the experiment. Females were mated following confirmation of pregestational body weight and glucose tolerance. For the discovery cohort, 15 LFt-HFb and 20 HFt-LFb females were used. Due to increased variability in body weight gain and glucose tolerance response that we observed in the discovery cohort, 20 LFt-HFb and 50 HFt-LFb females were used for the validation cohort.

### Longitudinal sampling and analysis of maternal fecal microbiota

To determine the impact of chronic depletion of soluble fiber on the background of a high-fat diet on gut microbiota prior and during pregnancy, a serial sampling strategy was used to examine the impact of dietary transition, chronic HFt-LFb diet consumption, and impact of HFt-LFb diet of pregnancy-associated gut microbiota dynamics. Fecal pellets were collected prior to transition to HFt-LFb diet, weekly fecal samples were collected during consumption of either HFt-LFb or LFt-HFb diet, and upon confirmation of copulation plug, daily fecal samples were collected during pregnancy.

### Cultivation and culture of *Gardnerella vaginalis* 11E4

*Gardnerella vaginalis* isolate 11E4 were maintained on Human Bilayer Tween Agar (BD) plates and NYC III medium according to the manufacturer’s instructions. Agar plates and liquid cultures were incubated at 37 °C with anaerobic gas mixture, 80% N2, 10% CO2, and 10% H2. Fresh live cultures were used for inoculation experiments and density was confirmed before each inoculation by dilution plating and counting colony forming units (CFU).

### Midgestational colonization by *Gardnerella vaginalis* strain 11E4 to model presence of a common member of CST IV

To model the local and peripheral inflammation observed in the presence of *G. vaginalis* in the cervicovaginal space of pregnant women in a clinical setting, pregnant mice were inoculated intravaginally with 50µL of 2 x 10^8^ CFU/ml saline of *G. vaginalis* strain 11E4 at gestational day 13.5 and E15.5, similar to previous reports (Gilbert et al., 2013; Sierra et al., 2018). *G. vaginalis* strain 11E4 was isolated from a women with BV (Nugent Score 7), a vaginal pH of 4.7, and a CST IV vaginal microbiota (subject 15 in (Ravel et al., 2013). The strain carries all known pathogenic traits associated with *G. vaginalis* proper, including the expression of vaginolysin, a cholesterol-dependent pore-forming cytolysins implicated in vaginal dysbiosis (Gelber et al., 2008; Ragaliauskas et al., 2019). This strain was selected based on pilot testing conducted by the Ravel lab in which 14 strains of human *G. vaginalis* were evaluated for biofilm formation, a feature associated with symptomatic bacterial vaginosis (Gelber et al., 2008; Ragaliauskas et al., 2019). Based on this, we tested two biofilm forming strains (11E4 and 16B1) and confirmed the presence of *G. vaginalis* 48-hours post-inoculation in mouse vaginal fluid by qPCR for *rpoB*. Days of inoculation (E13.5, E15.5) were selected based on previous work showing that introduction of *G. vaginalis* on these days induces an inflammatory response without significantly impacting placentation or inducing delivery. Live *G. vaginalis* cultures were inoculated at each time point, and colony forming unit counts were performed on Human Bilayer Tween agar for each batch of *G. vaginalis* cultures to ensure that females are inoculated with similar densities of pure live *G. vaginalis* cultures that were free of contamination.

### Genomic DNA isolation and 16S rRNA marker gene sequencing

Genomic DNA from fecal pellet samples were isolated using the Stratec PSP Spin Stool DNA Plus kit using the difficult to lyse bacteria protocol from the manufacturer (STRATEC Molecular GmbH, Berlin, Germany). Each sample DNA was eluted into 100 µL of Elution Buffer provided by the Stratec PSP Spin Stool DNA Plus kit. The V4 region of the bacterial 16S rRNA gene was amplified using a dual-index paired-end sequencing strategy for the Illumina platform as previously described^45^. Sequencing was performed on a MiSeq instrument (Illumina, San Diego, CA) using 2×250 base paired-end chemistry at the University of Maryland School of Medicine Institute for Genome Sciences. The sequences were demultiplexed using the dual-barcode strategy, a mapping file linking barcode to samples and split_libraries.py, a QIIME-dependent script. The resulting forward and reverse fastq files were split by sample using the QIIME-dependent script split_sequence_file_on_sample_ids.py, and primer sequences were removed using TagCleaner (version 0.16). Further processing followed the DADA2 workflow for Big Data and DADA2 (v.1.5.2) (https://benjjneb.github.io/dada2/bigdata.html). Data filtering was set to include features where 20% of its values contain a minimum of 4 counts. In addition, features that exhibit low variance across treatment conditions are unlikely to be associated with treatment conditions, and therefore variance was measured by inter-quartile range and removed at 10%. Data was normalized by cumulative sum scaling and differential abundance analysis was conducted using Linear Discriminant Analysis effect size with an FDR cut-off at q < 0.05. For quality control purposes, water and processed blank samples were sequenced and analyzed through the bioinformatics pipeline. Taxa identified as cyanobacteria or ‘unclassified’ to the phylum level were removed. For confirmation of human vaginal microbiota, taxonomic assignments were performed using a combination of a phylogenetics-based classifier and speciateIT software (http://www.speciateIT.sourceforge.net)

### Bulk RNA sequencing of fetal and postnatal tissues

On E18.5, pregnant dams were rapidly decapitated by cervical dislocation. Litter characteristics, such as intrauterine position, number of offspring, sex ratio, and resorption sites, were noted. Tail snip from embryos were collected and retained for determination of sex by genotyping using primers specific to Jarid1 (5′-TGAAGCTTTTGGCTTTGAG-3′ and 5′-CCGCTGCCAAATTCTTTGG-3′). Reactions were incubated at 94°C for 5 min, followed by 35 cycles of 94°C for 20 s, 54°C for 1 min, and 72°C for 40 s, followed by 72°C for 10 min. Following thermal cycling, 15 µL from each PCR product were mixed with 5 µL 6x loading dye solution (Sigma) before being loaded onto a 2% (w/v) agarose gel (Sigma) next to 0.5 µg 100-bp DNA ladder (Sigma) and electrophoresed in lx TAE buffer (40 mM Tris-acetate, 2 mM EDTA) at 120 V/cm for 45 min.

Fetal intestinal tracts were rapidly frozen in liquid nitrogen and maintained in –80 °C until RNA isolation. To control for the significant contribution of uterine horn laterality and intrauterine position, we selected fetal intestinal tract samples from conceptuses that were required to be from the first third of the embryos from cervical end; male samples flanked by two females (2F males) were excluded, and embryos could not exhibit overt signs of developmental delay (for example, small size relative to litters and no presence of a neighboring resorption site). Tissue was homogenized in Trizol, and messenger RNA was purified using the RNeasy Mini kit (Qiagen). Illumina single-end cDNA libraries of E18.5 fetal intestinal tract mRNA was prepared from 250 ng total RNA using the TruSeq Stranded mRNA Kit with poly-A enrichment (RS-122-2101, Illumina) according to the manufacturer’s protocol. Library fragment size was quantified using an Agilent High Sensitivity D1000 ScreenTape Assay on an Agilent 4200 Tapestation System (G2991AA). Library concentration was quantified using the Qubit dsDNA HS (High Sensitivity) Assay Kit. Samples representative of all treatment groups were multiplexed and sequenced on an Illumina NextSeq500 instrument using high-output 1×75-bp geometry (Illumina).

### Quantitative analysis of vaginal bacteria by qPCR

Genomic DNA from vaginal fluid recovered in 50 µL saline were isolated using the Stratec PSP Spin Stool DNA Plus kit using the difficult to lyse bacteria protocol from the manufacturer (STRATEC Molecular GmbH, Berlin, Germany). Each sample DNA was eluted into 100 µL of Elution Buffer provided by the Stratec PSP Spin Stool DNA Plus kit. Gene copy number of *G. vaginalis rpoB* was determined by quantitative real-time PCR analysis, performed on an Applied Biosystems QuantStudio6 Flex Real Time PCR System (Life Technologies, Grand Island, NY). Genomic DNA was extracted using Stratec PSP Spin Stool DNA Plus kit from vaginal swabs, TaqMan Assays specific for *G. vaginalis rpoB*, panbacterial 16S rRNA genes, TaqMan Vaginal Microbiota Amplification Control, and TaqMan Fast Advanced Master Mix (Life Technologies). Gene copy number counts were calculated using a standard curve method and the 16S rRNA gene level as an internal standard.

### RNA-seq analysis of the adult paraventricular nucleus of the hypothalamus (PVN)

Frozen brains from postnatal day 56 males were cryosectioned at −20 °C. Using a hollow 1.0 mm needle, the PVN was removed according to the mouse brain atlas (Paxinos and Franklin, 2019). PVN micropunches were immediately dispensed into 500 µl of Trizol and stored at −80 °C until processing. Messenger RNA was purified RNeasy Mini kit (Qiagen). Illumina single-end cDNA libraries of adult male PVN mRNA were prepared from 250 ng total RNA using the TruSeq Stranded mRNA Kit with poly-A enrichment (RS-122-2101, Illumina) according to the manufacturer’s protocol. Samples were multiplexed and sequenced on two identical NextSeq500 lanes using high-output 1×75-bp chemistry (Illumina). The concatenated FASTQ files generated from Illumina were used as input to kallisto, a program that pseudoaligns high-throughput sequencing reads to the *Mus musculus* reference transcriptome (version 38) and quantifies transcript expression. We used 60 bootstrap samples to ensure accurate transcript quantification. Gene isoforms were collapsed to gene symbols using the Bioconductor package tximport (version 3.4). Genes were filtered to counts per million > 1 in at least 3 samples. The filtered gene list was normalized using trimmed mean of M-values in edgeR. Variance weights were calculated using voom, and differential expression analysis of linear fit models was performed using limma v.3.12.3.

### Depletion of maternal microbiota

Pregnant dams were provided with ad libitum access to drinking water mixed with ampicillin, vancomycin and neomycin (all 1mg/ml), and supplemented with 1.125 g aspartame to increase palatability, starting on gestational day 15. Vehicle-treated females received aspartame in drinking water. The dams and the neonatal mice continued to receive antibiotic-containing drinking water until tissue collection at postnatal day 1.

### Blood lymphocyte counts

Blood (30 µl) was collected from postnatal day 1, 2, 3, 21 and 60 male offspring into EDTA-loaded tubes, and lymphocyte absolute counts and frequencies in the whole blood were determined using a Heska Element HT5 Veterinary Analyzer at the ZooQuatic Laboratory within the Institute of Marine and Environmental Technology.

### Whole blood sample collection and storage

Thirty microliters whole blood from postnatal day 1 and 100 µL whole blood from postnatal day 56 mice was collect for mass cytometry analysis and mixed with Cytodelics Whole Blood Cell Stabilizer at a 1:1 ratio, incubated at room temperature for 10 minutes and transferred to a −80°C freezer for long-term storage awaiting analysis. Whole blood samples preserved in Cytodelics stabilized were thawed at 20°C for 5 minutes. Samples were then diluted 1:4 with Cytodelics Wash Buffer #1 and allowed to lyse red blood cells for 15 minutes. Cells were then washed twice with Cytodelics Wash Buffer #2, filtered through a 35 micron mesh, cells were counted on a hemacytometer, and immediately proceeded with barcoding and staining.

### Antibody Labeling and Custom Conjugation

Purified monoclonal antibodies were either purchased pre-conjugated from Fluidigm or purchased from BioLegend in Maxpar Ready Purified Antibody formulation and conjugated in-house using the MAXPAR X8 polymer conjugation kit (Fluidigm Inc.) according to manufacturer’s protocol. Antibody concentration before and after conjugation was measured by NanoDrop, antibodies were diluted in Stabilizer, and antibodies were titrated prior to use in the staining panel.

### Mass Cytometry Barcoding, Staining and Acquisition

A maximum of 2×10^6^ cells per whole blood sample were barcoded using six combinations of palladium isotopes (^102^Pd, ^104^Pd, ^105^Pd, ^106^Pd, ^108^Pd, ^110^Pd) with the 20-Plex Pd barcoding kit (Fluidigm). Cells were fixed for 10 minutes in 1mL of Fix I Buffer at room temperature, followed by two washes in 1 mL of MaxPar Perm Buffer. Barcodes were resuspended in 100 µL Perm Buffer, transferred to each sample and incubated for 30 minutes at room temperatures. All samples were then washed twice with Cell Staining Buffer (CSB) and then pooled. Cells were suspended in 150ml antibody cocktail per 10^7^ cells and incubated for 30 minutes at room temperature. Antibodies used are listed in **Key Resources Table.** After staining, cells were washed twice with CSB followed by overnight incubation in Iridium-labeled DNA-intercalator in 4% formaldehyde for a final concentration of 0.125 mM at 4°C. Following two washes in Cell Staining Buffer, cells were rapidly frozen cryoprotective medium consisting of 90% FBS/10% DMSO. Following freezing, samples were shipped overnight to the University of Virginia School of Medicine Flow Cytometry Facility. On the day of acquisition, cells were washed once in Call Acquisition Buffer, once in PBS and twice in milliQ H2O filtered through a 35mm nylon mesh and counted. Cells were diluted in milliQ H2O containing 10% EQ Four Element Calibration Beads. Samples were acquired a CyTOF2 mass cytometers with a wide bore configuration, using noise reduction, event length limits of 10-150 pushes and a sigma value of 3. Cells were acquired at a flow rate of 0.045ml/min.

### Mass Cytometry Preprocessing and Gating

All FCS files were preprocessed using Gaussian discrimination parameters, following recommendations by Fluidigm. Following this preprocessing step, FCS files were uploaded to the Astrolabe Cytometry Platform (Astrolabe Diagnostics, Inc.) where transformation, debarcoding, cleaning, labeling, and unsupervised clustering was done. Data was transformed using arcsinh with a cofactor of 5. Experimental batches were debarcoded and individual samples were then labeled using the Ek’Balam algorithm, a hierarchy-based algorithm for labeling cell subsets which combines knowledge-based gating strategy with unsupervised clustering using the FlowSOM algorithm. Differential expression analysis to compare treatment effects was conducted within the Astrolabe Cytometry Platform. Groups average t-SNE maps and unsupervised clustering using FlowSOM were generated in Cytobank.

## Quantification and Statistical Analysis

Statistical information including sample size, mean, and statistical significance values are indicated in the text or the figure legends. A variety of statistical analyses were applied, each one specifically appropriate for the data and hypothesis, using the R statistical environment. For standard metabolic endpoints, analysis of variance (ANOVA) testing with repeated-measures corrections and Tukey post-hoc tests were utilized, with significance at an adjusted p < 0.05. Processing of RNA-Seq data were conducted using standardized and published protocols. Cytobank and Astrolabe Diagnostics were used for analysis of CyTOF data. No custom script was used to analyze RNA sequencing or cytometric data.

---

**Supporting Figure 1.**
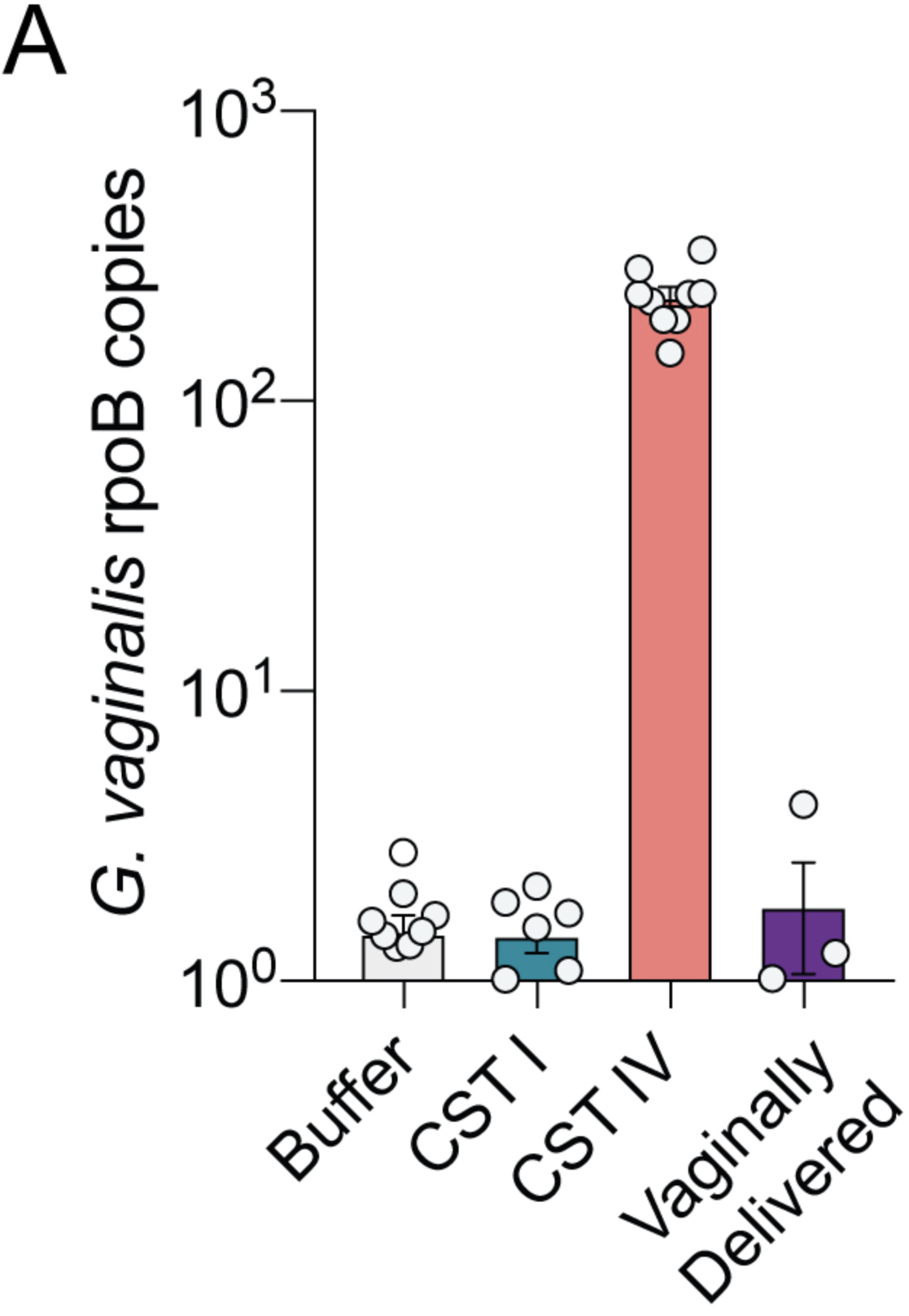
Quantitative PCR-based validation of human vaginal microbiota in the intestine of C-section delivered mice exposed to Amies, CST I or CST IV. C-section mice inoculated with CST IV show increased rpoB copy number compared with Amies, CST I and vaginally delivered offspring. Copy number of *G. vaginalis* was quantified using TaqMan assay for *rpoB*, and the TaqMan vaginal microbiota amplification control was used to evaluate amplification efficiency. N = 8 Amies, 6 CST I, 9 CST IV, 3 vaginally delivered. Data represented as mean ± SEM with individual data points overlaid.

**Supporting Figure 2.**
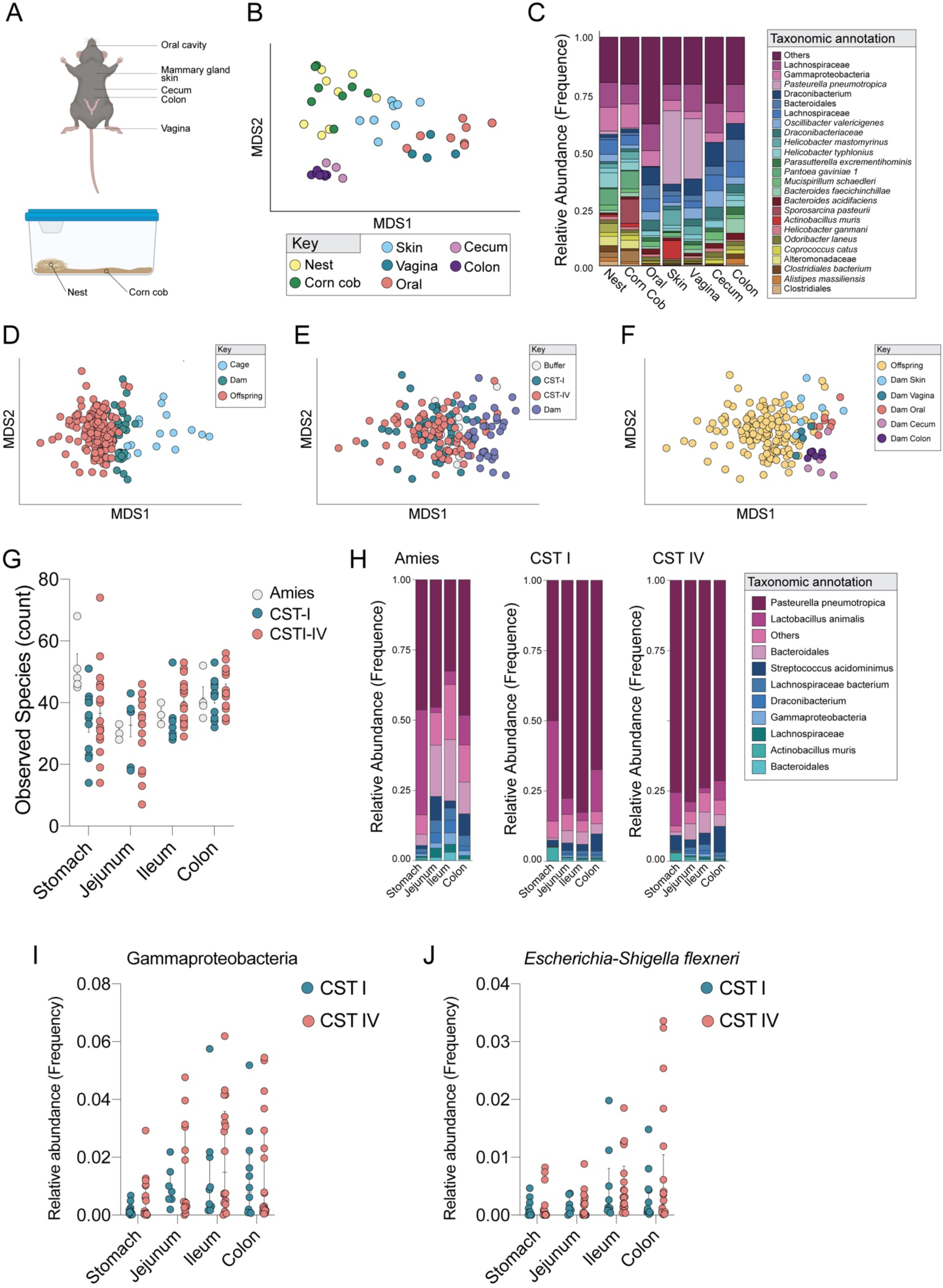
Defining the environmental and maternal microbial reservoirs involved in vertical transmission of microbiota from foster dam to C-section offspring. **(A)** Schematic of the samples collected to determine the microbial reservoirs involved in vertical transmission. The local cage environment was assessed by sampling nestlet material on which pups had direct physical contact, and corn cob was sampled surrounded the nestlet. Maternal samples were collected as depicted in the schematic. **(B)** NMDS analysis of environmental and maternal samples showing that clustering was influenced by maternal body site. **(C)** Mean relative abundance of the top 20 taxa across maternal body sites and the local cage environment, demonstrating a bloom of *P. pneumotropica* on the mammary gland associated skin and vagina of the foster dam. **(D)** NMDS analysis of offspring, foster dam, and cage samples showing distinct clustering of offspring and cage samples, with foster dam as an intermediary. N = 3 – 20 samples per treatment and intestinal segment; 15 cage samples; 32 foster dam samples. **(E)** NMDS analysis showing no distinct clustering between gut samples from Amies, CST I and CST IV inoculated C-section pups. N = 128 C-section pup samples; 32 foster dam samples. **(F)** NMDS analysis showing closest clustering between C-section pup gut samples and the foster dams compared with samples from the vagina, cecum, colon and oral cavity of the foster dam. N = 128 C-section pup samples; 32 foster dam samples. **(G)** Barplot demonstrating intestinal region specific differences in alpha diversity in Amies, CST I, and CST IV C-section pups. N = 3 – 20 samples per treatment and intestinal segment. Data represented as mean ± SEM with individual data points overlaid. **(H)** Mean relative abundance of the top 20 taxa across intestinal segments between Amies, CST I, CST IV C-section pups demonstrating that *P. pneumotropica* as a dominant taxa in the neonate gut. N = 3 – 20 samples per treatment and intestinal segment. **(I)** LEfSe analysis demonstrating increased abundance of Gammproteobacteria in CST IV inoculated C-section offspring in an intestinal region-specific manner (FDR = 0.007, LDAscore = 4.86). N = 3 – 20 samples per treatment and intestinal segment. Data represented as median ± IQR with individual data points overlaid. **(J)** LEfSe analysis demonstrating increased abundance of Escherichia-Shigella in CST IV inoculated C-section offspring in an intestinal region-specific manner (FDR = 0.0003, LDAscore = 4.34). N = 3 – 20 samples per treatment and intestinal segment. Data represented as median ± IQR with individual data points overlaid.

**Supporting Figure 3.**
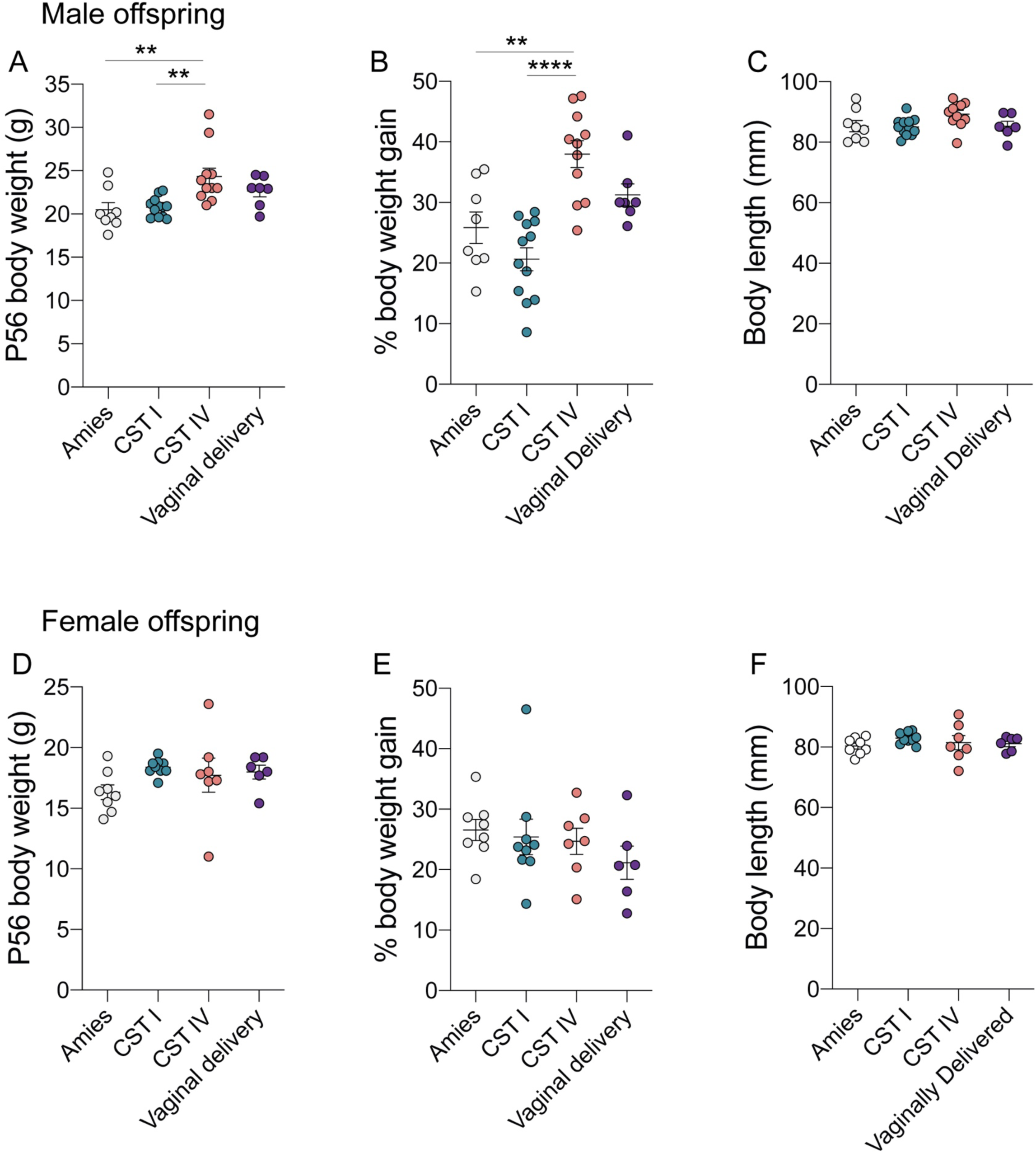
Exposure to human cervicovaginal vaginal microbiota impacts body weight across development in a sex-specific manner. **(A)** CST IV males weighed more than CST I and Amies males (one-way ANOVA, main effect of treatment, F_3, 34_ = 6.280, P=0.0017); Tukey’s post-hoc, CST I vs. CST IV, *P* = 0.0039; Amies vs. CST IV, *P* = 0.0050). N= 7 – 12 males per group. Data represented as mean ± SEM with individual data points overlaid. ** *P <* 0.01. **(B)** CST IV males showed increased percent body weight gain from P28 to P56 compared with CST I and Amies males F_3, 34_ = 13.58, *P* < 0.0001; Tukey’s post-hoc, CST I vs. CST IV, *P* < 0.0001; Amies vs. CST IV, *P* = 0.0024). N= 7 – 12 males per group. Data represented as mean ± SEM with individual data points overlaid. ** *P <* 0.01, *** *P <* 0.001. **(C)** Body weight differences were not due to differences in body length between Amies, CST I, CST IV, and vaginally delivered males at P56. N= 7 – 12 males per group. Data represented as mean ± SEM with individual data points overlaid. **(D-F)** No differences in P56 body weight, percent body weight gain, or body length were observed between Amies, CST I, CST IV, and vaginally delivered female offspring. N= 6-9 females per group. Data represented as mean ± SEM with individual data points overlaid. Data represented as mean ± SEM with individual data points overlaid.

**Supporting Figure 4.**
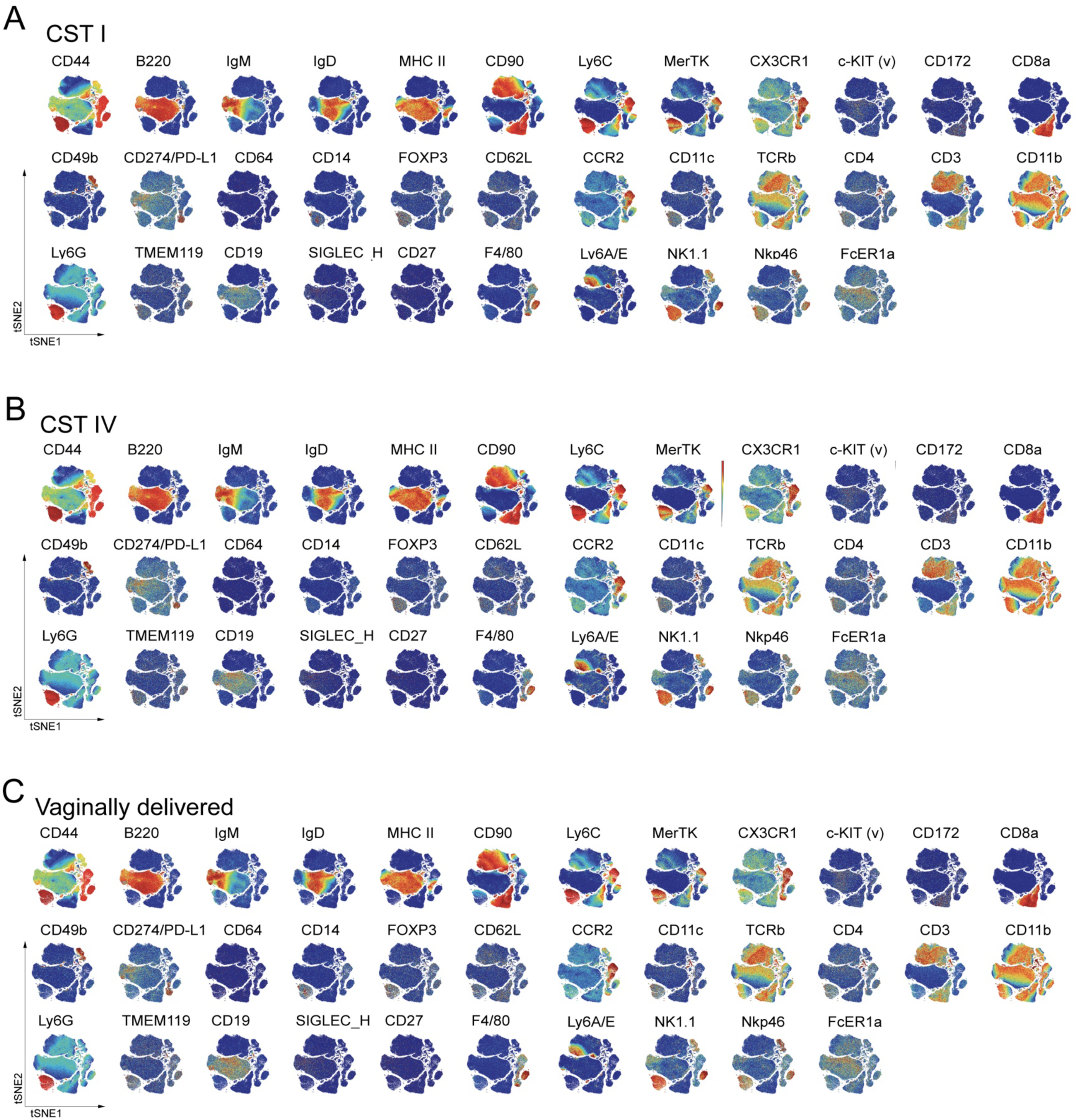
High-dimensional analysis of mass cytometry data using viSNE comparing the lasting effects of mode of delivery and colonization by human vaginal microbiota on the circulating immune compartment of cesarean delivered males inoculated with (A) CST I or (B) CST IV and (C) vaginally delivered males at postnatal day 56. viSNE maps were generated using equal sampling across treatment groups and the markers shown were used to generate the maps (n= 390,000 total events sampled, representative average from N = 3 −7 males per treatment).

**Supporting Figure 5.**
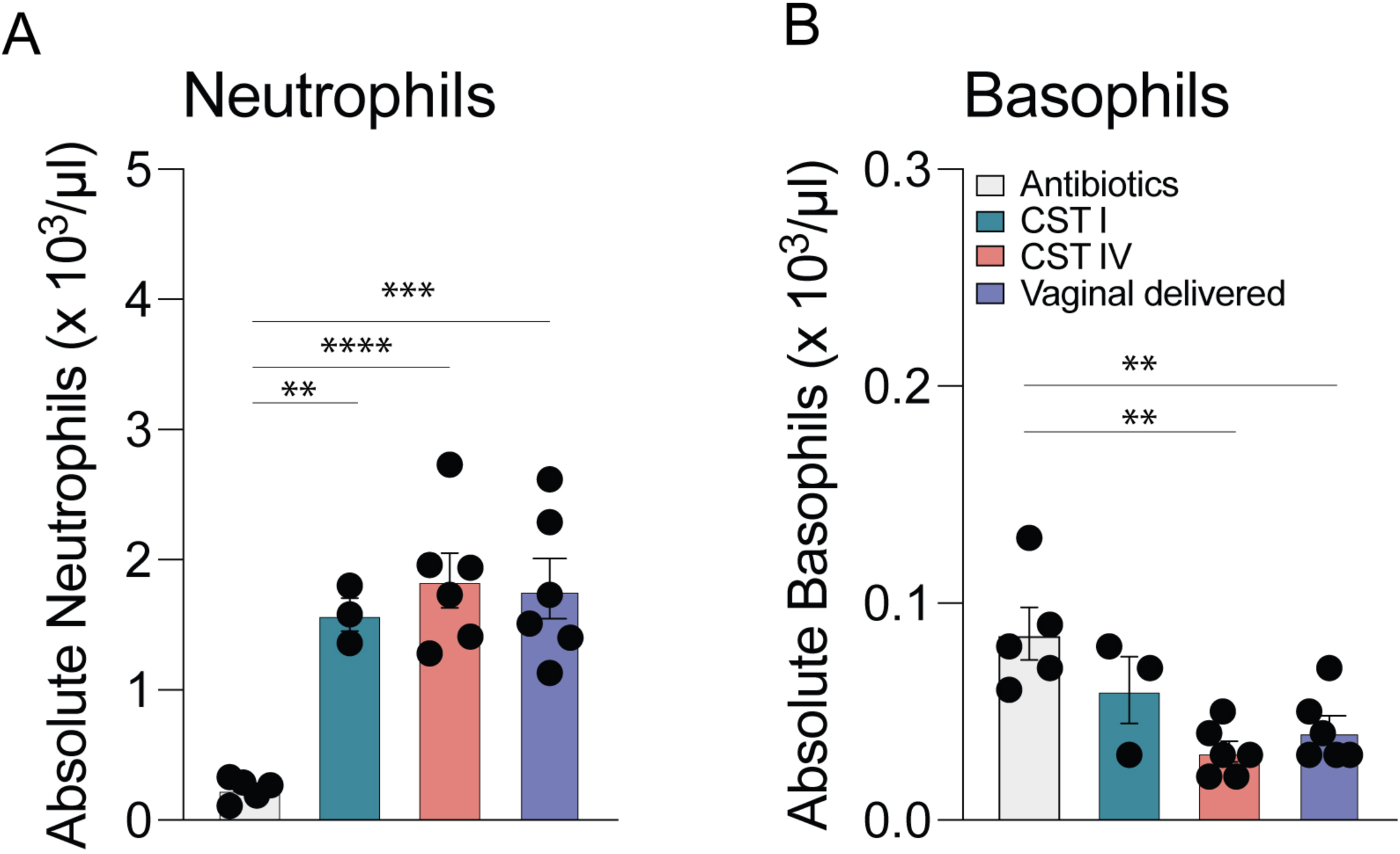
Early life colonization influences number of circulating neutrophils and basophils in males 24 hours following inoculation with human cervicovaginal microbiota. To determine whether early life colonization is necessary for expansion of neutrophil counts in circulation, we exposed pregnant dams to a clinically relevant combination of ampicillin, gentamicin and vancomycin (all 1 mg ml^−1^) starting at gestational day 15 to postnatal day 1. On postnatal day 1, corresponding to 24 hr post-inoculation, absolute white blood cell counts were quantified on an Element HT5 veterinary hematology analyzer. **(A)** Neutrophil count is significantly decreased in P1 antibiotic-exposed males compared with CST I, CST IV and vaginally delivered males (one-way ANOVA, main effect of treatment, F_3, 16_ = 15.46, *P <* 0.0001; Tukey’s post-hoc, Antibiotics vs. CST I, *P* = 0.0034; Antibiotics vs. CST IV *P* <0.0001; P2 Antibiotics vs. VD *P* = 0.0001). N = 3 – 6 males per treatment. Data are represented as mean ± SEM. ** *P <* 0.01, *** *P <* 0.001, **** *P <* 0.0001. **(B)** Basophil count is significantly increased in P1 antibiotic-exposed males compared with CST IV and vaginally delivered males (one-way ANOVA, main effect of treatment, F (3, 16) = 7.710, P=0.002; Tukey’s post-hoc, Antibiotics vs. CST IV *P* = 0.0018; P2 Antibiotics vs. VD *P* = 0.0096). N = 3 – 6 males per treatment. Data are represented as mean ± SEM. ** *P <* 0.01.

**Supporting Figure 6.**
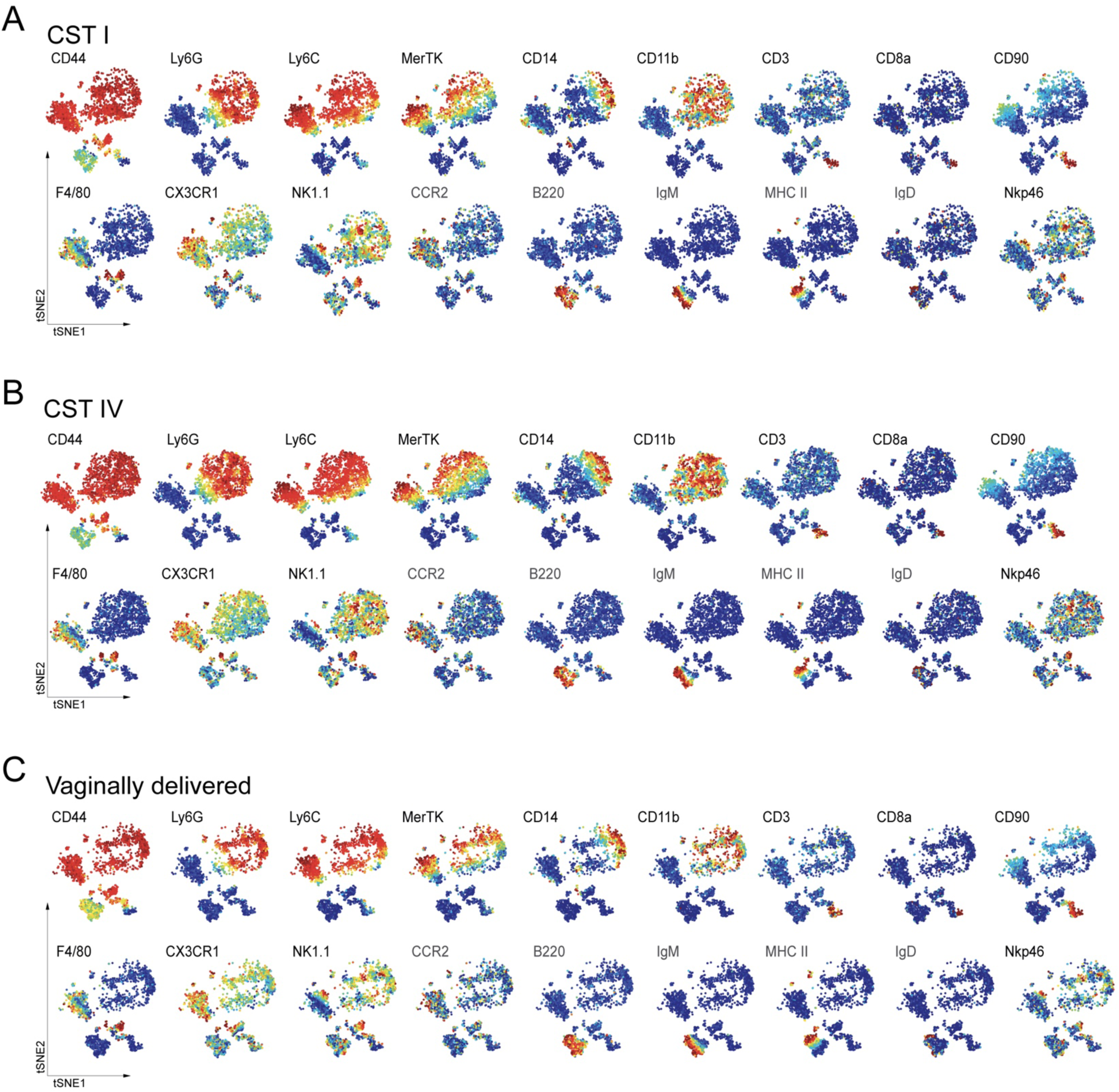
High-dimensional analysis of mass cytometry data using viSNE comparing the circulating immune compartment of cesarean delivered males inoculated with human vaginal (A) CST I or (B) CST IV and (C) vaginally delivered males at postnatal day 1, corresponding to 24 hrs post-inoculation. viSNE maps were generated using equal sampling and the markers shown were used to generate the maps (n = 10,000 total events sampled, representative average from N = 3 males per treatment).

**Supporting Figure 7.**
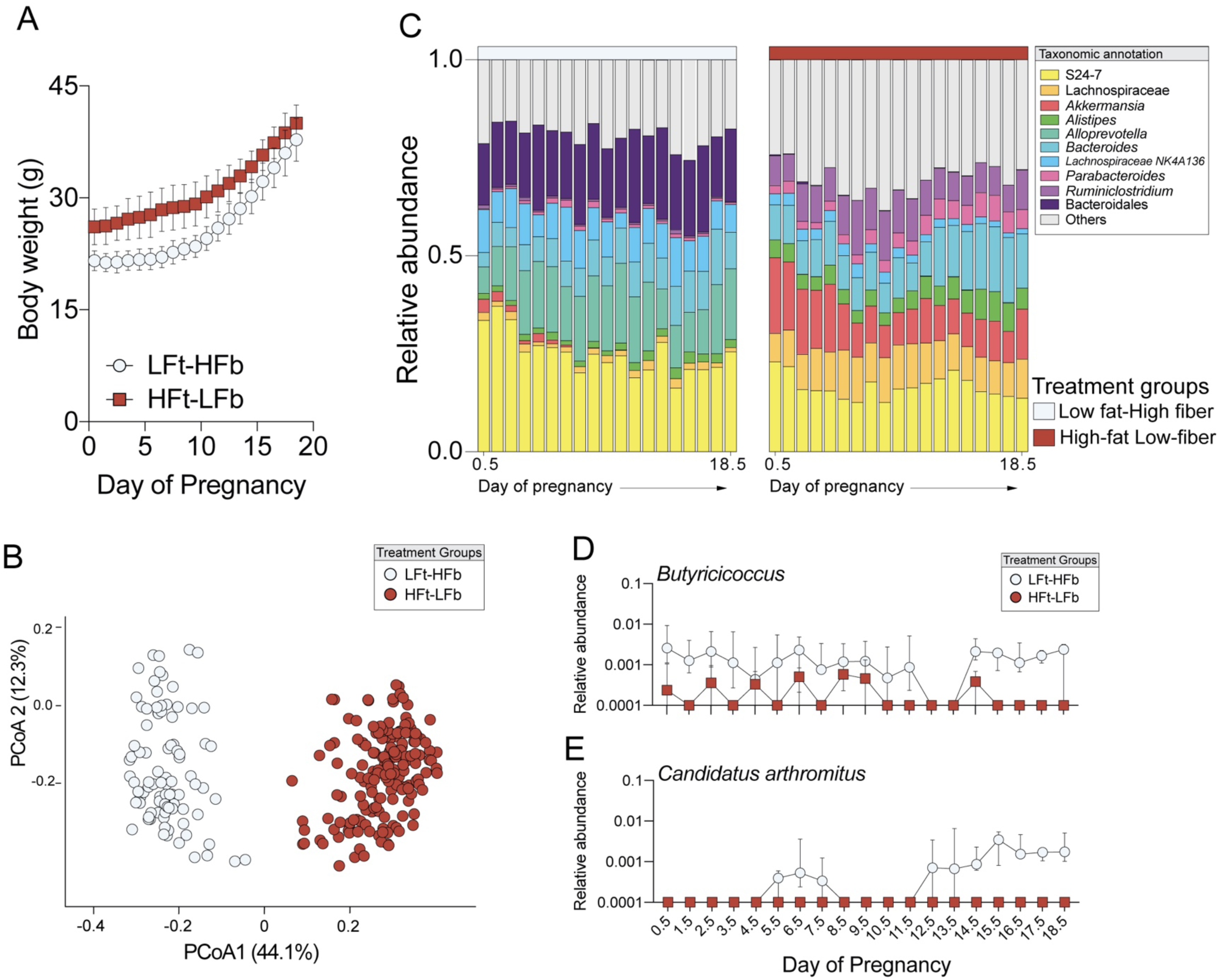
Maternal consumption of a high fat-low fiber diet results in persistent disruption to body weight and gut microbiota structure and composition during pregnancy. **(A)** Excessive maternal weight gain during pregnancy in dams consuming a high-fat low-fiber diet (two-way ANOVA, main effect of time, F_18, 324_ = 474.24, *P* < 0.0001; main effect of diet, F_1, 18_ = 23.51, *P* < 0.0001; time*diet interaction, F_18, 324_ = 6.73, *P* < 0.0001). No effect of *G. vaginalis* 11E4 on body weight was observed, thus samples are shown collapsed by diet. N = 12 – 20 females per treatment and timepoint. Data represented as mean ± SEM with individual datapoints overlaid. Data is representative of two independent experiments **(B)** Principal coordinates analysis demonstrating distinct clustering between dams consuming a high-fat low fiber and low-fat high-fiber diet across pregnancy. No effect of *G. vaginalis* 11E4 on gut microbiota structure was observed, thus samples are shown collapsed by diet. N = 12 – 20 females per treatment and timepoint, total of 259 samples. **(C)** Mean relative abundance of top ten lasting disruption to the fecal microbiota during pregnancy in females consuming a high-fat low-fat diet. No effect of *G. vaginalis* 11E4 on gut microbiota composition was observed, thus samples are shown collapsed by diet. N = 12 – 20 females per treatment and timepoint, total of 259 samples. **(D)** LEfSe analysis demonstrating decreased relative abundance of the butyrate-producing taxa *Butyricoccus* in females consuming a high-fat low-fat diet (FDR =, LDAscore =). No effect of *G. vaginalis* 11E4 on *Butyricoccus* relative abundance was observed, thus samples are shown collapsed by diet. N = 12 – 20 females per treatment and timepoint, total of 259 samples. **(E)** LEfSe analysis demonstrating decreased abundance of the immunomodulatory *Candidatus Arthromitus* in females consuming a high-fat low-fat diet during late pregnancy (FDR =, LDAscore =). No effect of *G. vaginalis* 11E4 on *Candidatus Arthromitus* relative abundance was observed, thus samples are shown collapsed by diet. N = 12 – 20 females per treatment and timepoint, total of 259 samples.

**Supporting Figure 8.**
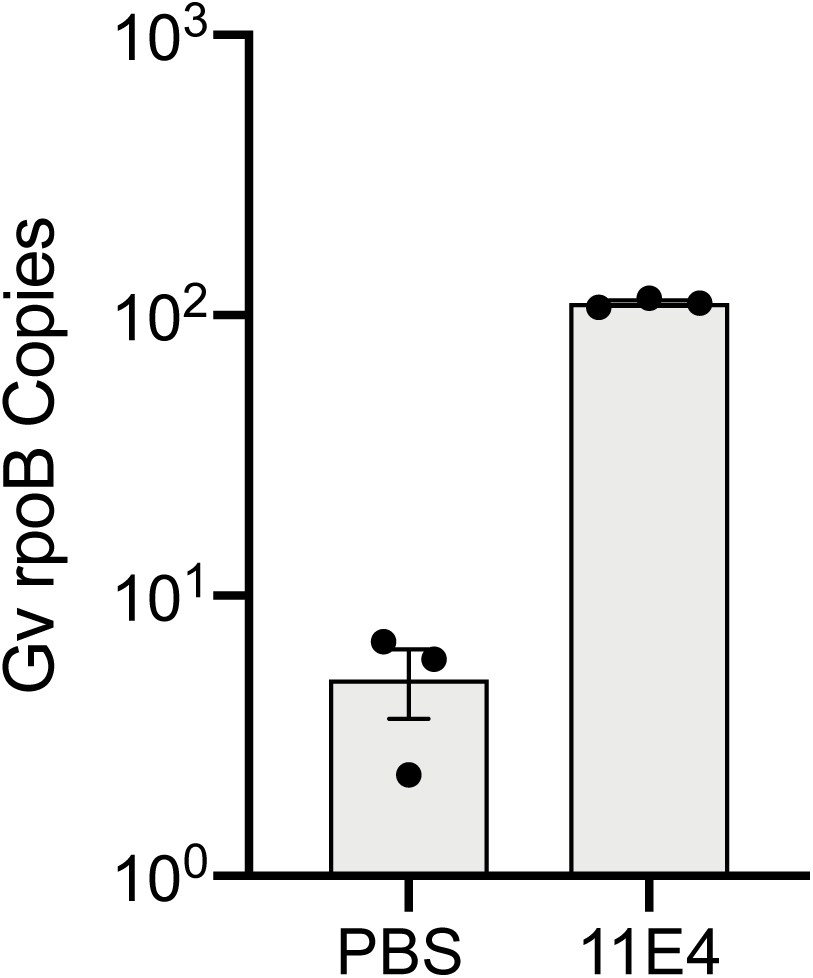
Confirmation of *G. vaginalis* isolate 11E4 in vaginal fluid 48hrs post-colonization. Presence of Gv was significantly increased in the vaginal fluid from Gv inoculated dams relative to PBS inoculated dams (Unpaired t-Test, t_4_ = 36.45, *P* < 0.0001). Data represented as mean ± SEM with individual data points overlaid.

**Supporting Figure 9.**
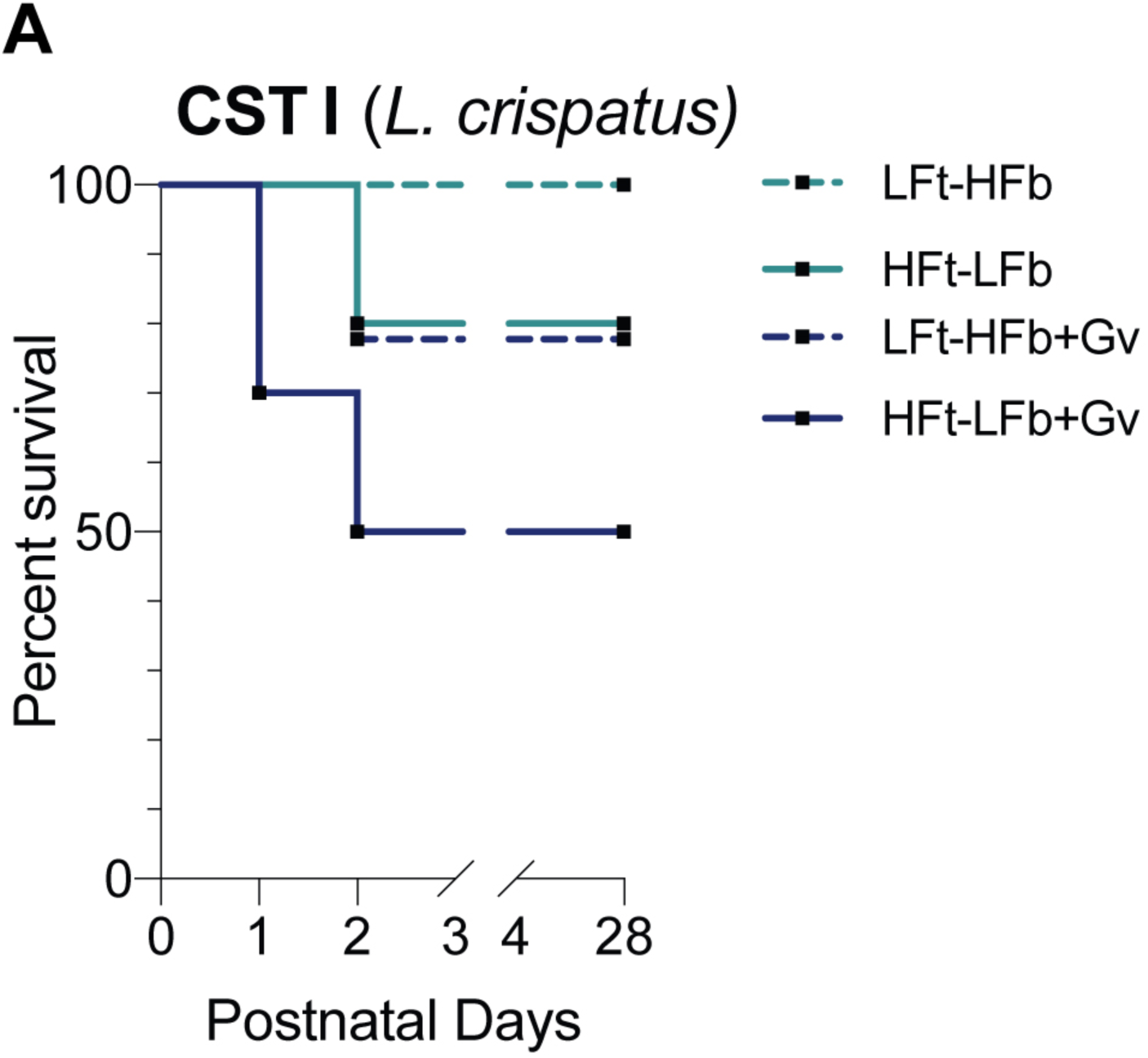
Combined effects of exposure to compounding maternal insults and colonization by human cervicovaginal CST I on offspring mortality. **(A)** Survival of offspring from dams that experience a single or multiple compounding adversities. All pups were c-section delivered and gavaged with human CST I inoculant. LFt-HFb = low-fat high-fiber; HFt-LFb = high-fat low-fiber; Gv = G. vaginalis 11E4. Kaplan-Meier survival analysis. N=15 offspring per treatment condition.

**Supporting Figure 10.**
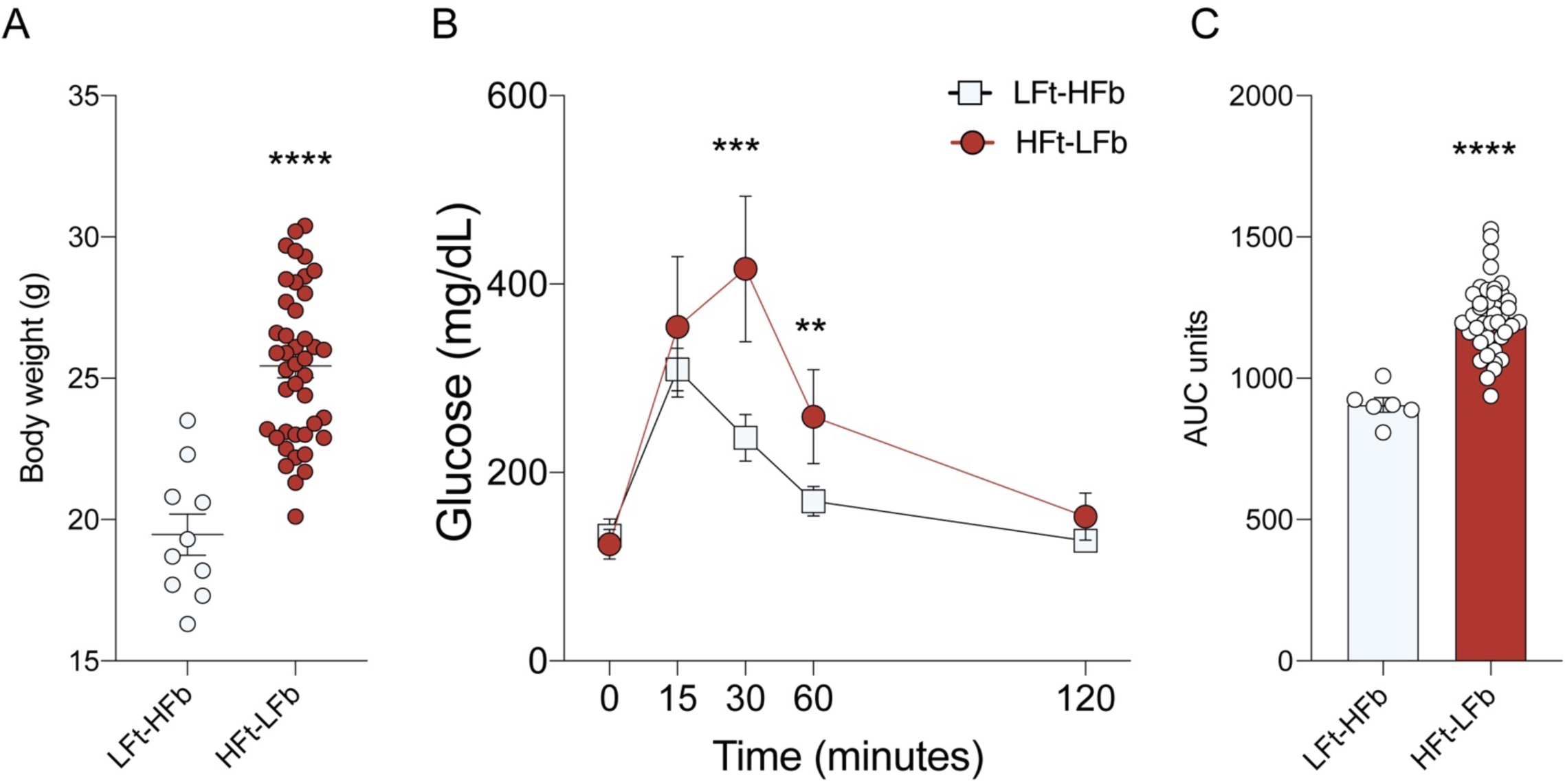
Validation of excessive maternal weight gain and glucose intolerance in females consuming a high fat-low fiber diet for six weeks. **(A)** Consumption of a high-fat low-fiber diet for six weeks increases body weight in females consuming a high-fat low-fiber diet compared with females consuming a low-fat high-fiber diet (two-tailed t-Test, *t_50_* = 6.408, *P* < 0.0001). N = 10 – 40 females per treatment. Data represented as mean ± SD. **** *P <* 0.0001. **(B)** Plasma levels of glucose levels during a glucose tolerance test in females consuming either a high-fat low-fiber or low-fat high-fiber diet. Females consuming a high-fat low-fiber diet showed significant delay in glucose clearance (two-way ANOVA, main effect of time, F_4, 180_ = 84.09, *P* < 0.0001; main effect of diet, F_1, 45_ = 29.65, *P* < 0.0001; time*diet interaction, F_4, 180_ = 11.56, *P* < 0.0001). ** *P <* 0.01; *** *P* < 0.001. **(C)** AUC of total plasma glucose levels showing increase glucose levels in females consuming a high-fat low-fiber diet (two-tailed T test, *t*_44_ = 5.821, *P <* 0.0001). Data represented as mean ± SD with individual datapoints overlaid. *** *P <* 0.001.

**Supporting Figure 11.**
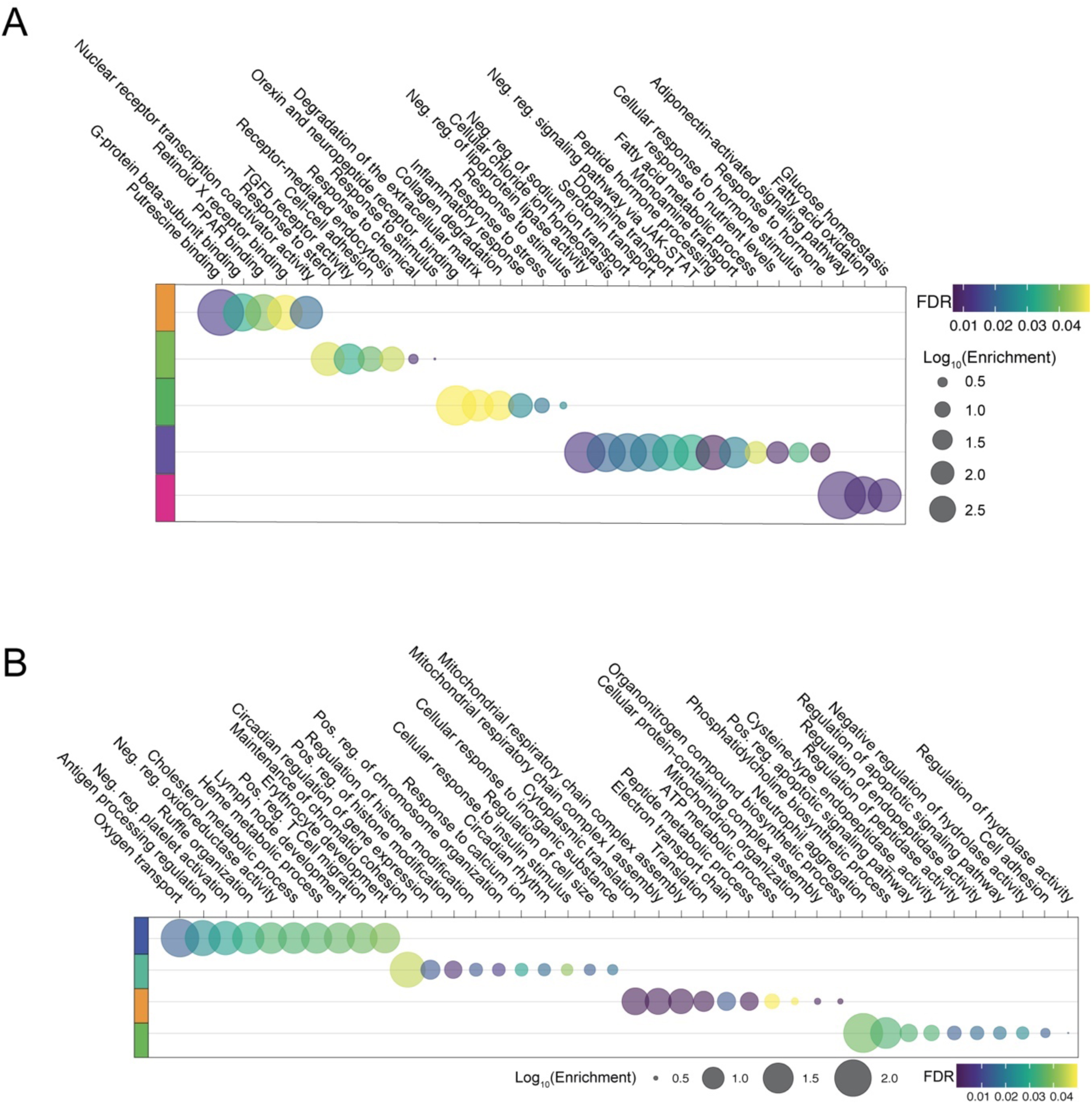
Cluster-based functional enrichment analysis showing significant effect of HFt-LFb and *G. vaginalis* and their combination on metabolic and immune pathways in the E18.5 placenta. **(A)** Cluster-based functional enrichment analysis of differentially expressed genes in the placenta of embryonic day 18.5 males, showing significant disruption in pathways involved in nutrient transport and fatty acid metabolism in the placenta of males exposed to a maternal high-fat low fiber diet and *G. vaginalis* vaginal colonization (FDR < 0.05). Bubble plot size denotes enrichment. N = 3 males per treatment. **(B)** Cluster-based functional enrichment analysis of differentially expressed genes in the ileum of embryonic day 18.5 males, showing significant disruption in pathways involved in tissue development, chromatin modification, immunity, and mitochondrial function in the ileum of males exposed to a maternal high-fat low fiber diet and G. vaginalis vaginal colonization (FDR < 0.05). Bubble plot size denotes enrichment. N = 3 males per treatment.

**Supporting Figure 12.**
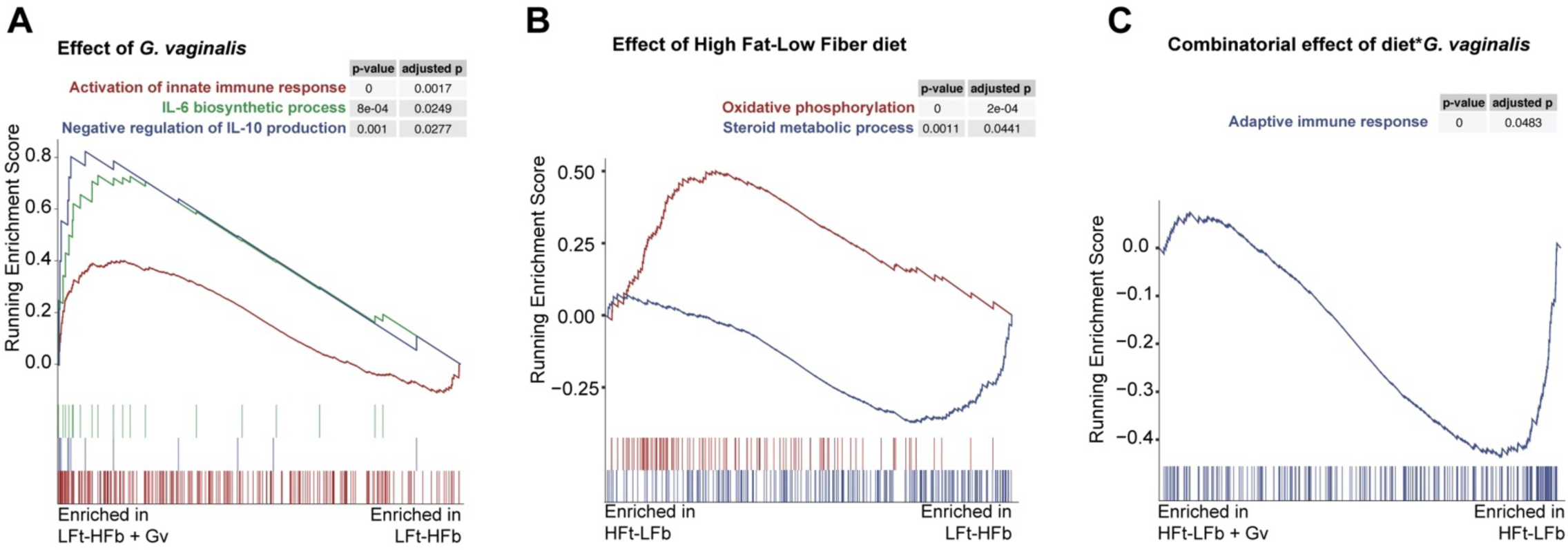
Gene set enrichment analysis (GSEA) showing significant effect of HFt-LFb and *G. vaginalis* and their combination on metabolic and immune pathways in the E18.5 placenta. **(A)** Analysis comparing the effects of *G. vaginalis* in LFt-HFb placentas shows significant enrichment of gene sets involved in activation of innate immune response, IL-6 biosynthesis, and negative regulation of IL-10 production relative to LFt-HFb placentas (N = 3 males per group, GSEA cut off FDR ≤ 0.05, NES > |1.5|). **(B)** Analysis comparing the effects of diet shows significant dysregulation of gene sets involved in oxidative phosphorylation and steroid metabolic processes in HFt-LFb male placentas relative to LFt-HFb male placentas (N = 3 males per group, GSEA cut off FDR ≤ 0.05, NES > |1.5|). **(C)** Analysis comparing the interaction between diet and *G. vaginalis* shows significant dysregulation of gene sets involved in the adaptive immune response in HFt-LFb + Gv male placentas relative to LFt-HFb male placentas (N = 3 males per group, GSEA cut off FDR ≤ 0.05, NES > |1.5|).

**Supporting Figure 13.**
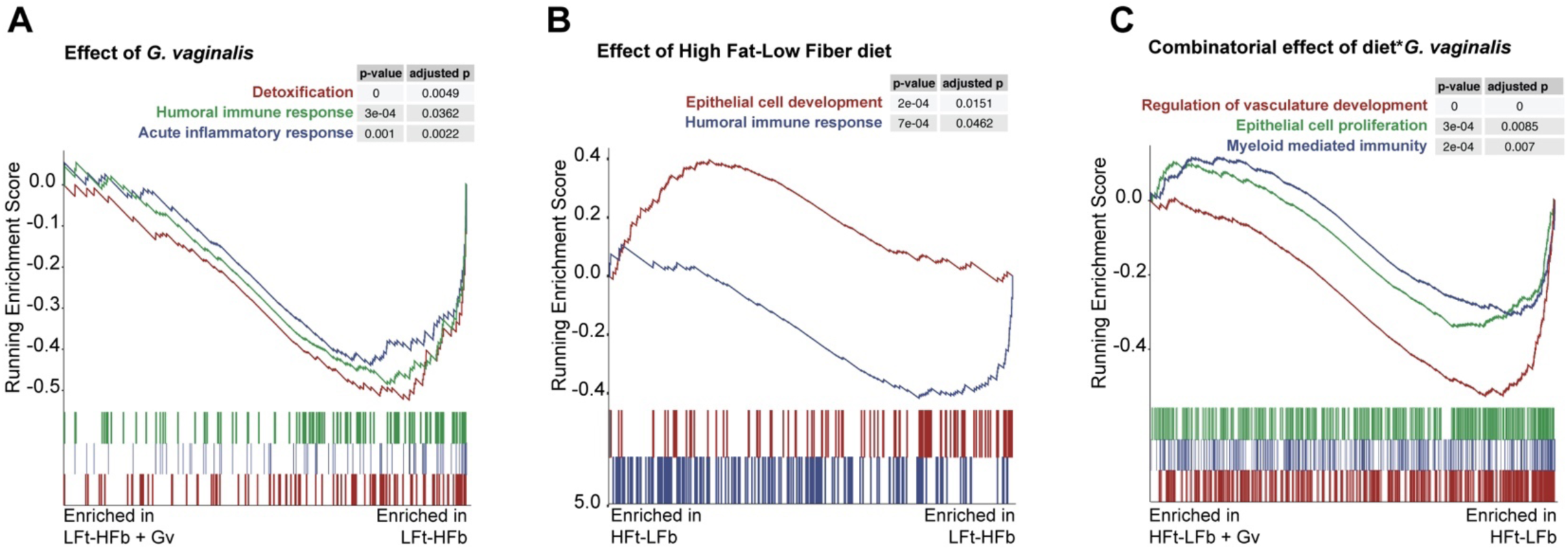
Gene set enrichment analysis (GSEA) showing significant effect of HFt-LFb and *G. vaginalis* and their combination on pathways involved in immune, epithelial and vasculature maturation of the E18.5 ileum. **(A)** Analysis comparing the effects of *G. vaginalis* in LFt-HFb ileum shows disruption in the gene sets involved in metabolic detoxification, the humoral immune response and the acute inflammatory response relative to the ileum of LFt-HFb pups (N = 3 males per group, GSEA cut off FDR ≤ 0.05, NES > |1.5|). **(B)** Analysis comparing the effects of diet shows significant dysregulation of gene sets involved in humoral immune response and epithelial cell development in HFt-LFb male ileum samples relative to LFt-HFb male ileum samples (N = 3 males per group, GSEA cut off FDR ≤ 0.05, NES > |1.5|). **(C)** Analysis comparing the interaction between diet and *G. vaginalis* shows significant dysregulation of gene sets involved in the regulation of vasculature development, epithelial cell proliferation and myeloid mediated immunity in HFt-LFb + Gv male ileum samples relative to LFt-HFb male ileum samples (N = 3 males per group, GSEA cut off FDR ≤ 0.05, NES > |1.5|).

